# Development of a Data-driven Integrative Model of Bacterial Chromosome

**DOI:** 10.1101/2023.01.29.526099

**Authors:** Abdul Wasim, Palash Bera, Jagannath Mondal

## Abstract

The chromosome of archetypal bacteria *E. coli* is known for a complex topology with 4.6 × 10^6^ base pairs (bp) long sequence of nucleotide packed within a micrometer-sized celllular confinement. The inherent organization underlying this chromosome eludes general consensus due to the lack of a high-resolution picture of its conformation. Here we present our development of an integrative model of *E. coli* at a 500 bp resolution (https://github.com/JMLab-tifrh/ecoli_finer), which optimally combines a set of multi-resolution genome-wide experimentally measured data within a framework of polymer based architecture. In particular the model is informed with intra-genome contact probability map at 5000 bp resolution derived via Hi-C experiment and RNA-sequencing data at 500 bp resolution. Via dynamical simulations, this data-driven polymer based model generates appropriate conformational ensemble commensurate with chromosome architectures that *E. coli* adopts. As a key hallmark, the model chromosome spontaneously self-organizes into a set of non-overlapping macrodomains and suitably locates plectonemic loops near the cell membrane. As novel extensions, it predicts a contact probability map simulated at a higher resolution than precedent experiments and can demonstrate segregation of chromosomes in a partially replicating cell. Finally, the modular nature of the model helps us to devise control simulations to quantify the individual role of key features in hierarchical organization of the bacterial chromosome.

## 1 Introduction

The *E. coli* chromosome though being only 4.64×10^6^ base pairs (bp) long^1^ still eludes us with its actual chromosome structure and a complete understanding of its organization. Though initially thought to be just a disorganized blob of DNA, it is now known to contain a hierarchically complex organization. The DNA, if stretched out, would measure ∼1.5 mm in length.^2^ It undergoes almost 1000 times compaction when packed inside a cell of length ∼2-3 *µm*. It has been observed that due to the efficient packing of chromosome inside the bacterial cell it occupies only 10-20% of its cell volume.^3^ Out of the mechanisms responsible for DNA compaction, DNA supercoiling ^4^ via plectoneme formation and DNA condensation^5–9^ via intra-chromosome contacts are two of the most important ones. Examples of other mechanisms that augment DNA compaction are the confinement effect due to the cell wall^10^ and macromolecular crowding from proteins, ribosomes and RNA.^11,12^

Over the past decade the emergence of chromosome conformation capture techniques^13–17^ and observations of its organization via microscopy^18,19^ have provided us with numerous glimpses into its organization in-vivo. One of the foremost features is its division into four macrodomains (Ori, Ter, Right, Left) and two non-structured regions (Non Structured Right (NSR) and Non Structured Left (NSL)).^20^ Ori (origin) contains the loci oriC from which chromosome replication initiates while Ter (terminus) contains the dif loci which is the last region of the chromosome to be replicated. The macrodomains were shown to be mutually non-overlapping, rather the DNA belonging to one macrodomain tend to interact more within themselves.^20^ The spatial orientation of the macrodomains were also seen to be dependent on the cell growth stage^21^ and a dynamical spatial orientation was observed based on the location of oriC and dif.^21^

Apart from intra-genome contact probability as predicted by chromosome conformation capture techniques (Figure 1a), other key mechanism by which the chromosome is condensed is via formation of dense, supercoiled structures called plectonemes (Figure 1b). Multiple proteins are responsible for supercoiling the DNA to form plectonemes. It is also a mechanism by which the bacteria controls expression of various genes encoded by its chromosome. Supercoiled regions cannot be read by RNAP, which is responsible for reading DNA to generate mRNA. Thus genes belonging to plectonemic DNA are excluded from being transcribed. On the other hand there are regions of the chromosome which undergo continuous transcription, such as operons, which are kept decondensed and plectoneme free for the RNAP. Experiments such as RNAP-ChIP^22^ or RNA-Seq^23^ (Figure 1c) can thus determine which regions of the DNA are being transcribed. This information can be then used to determined which sections of the chromosome have plectonemes (not transcribing) and which regions are plectoneme free (being transcribed) with near base pair resolutions.

**Figure 1:**
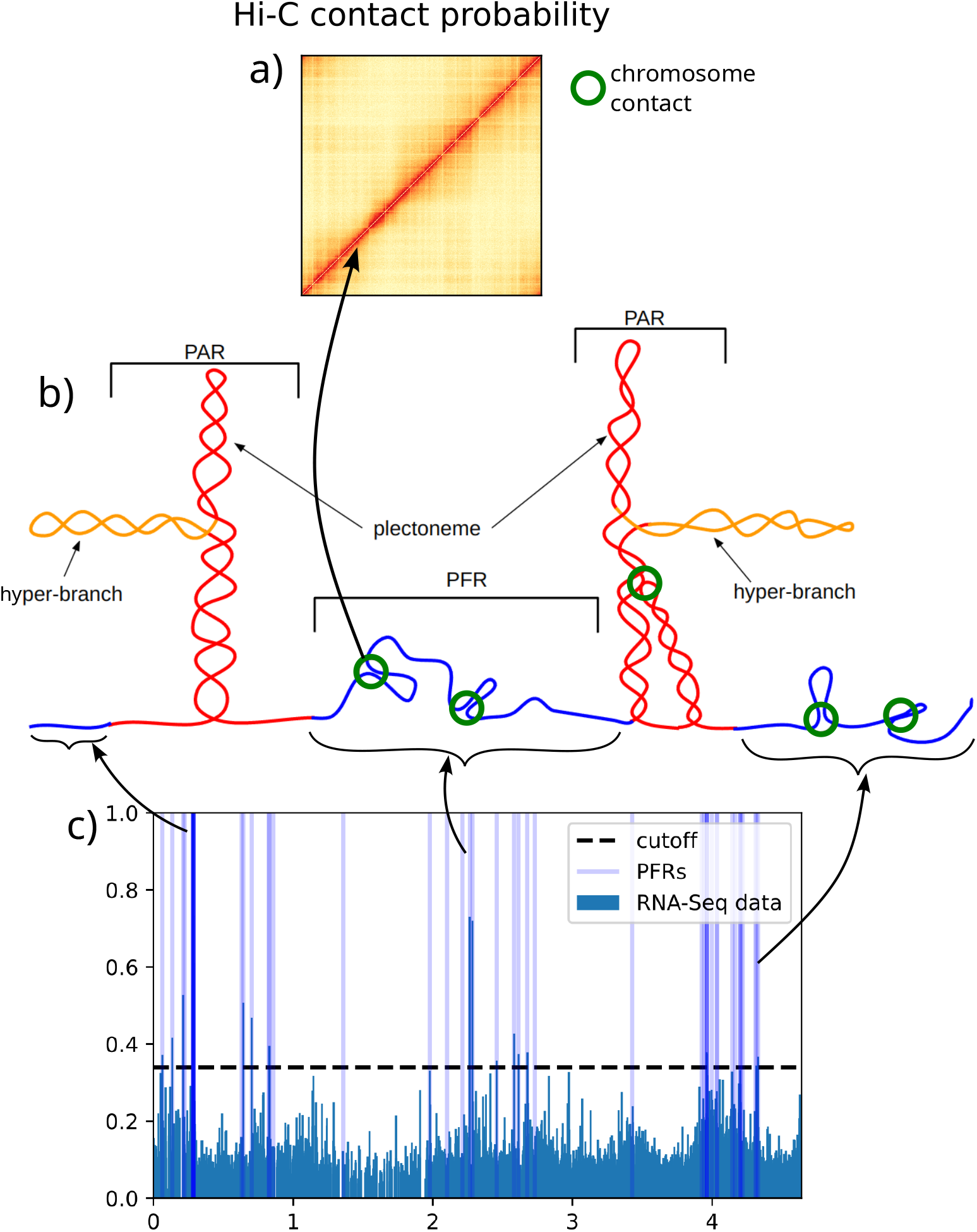
**a)** Hi-C contact probability matrix at 5000 bp. **b)**Cartoon representation of plectonemes. The green circles show-case chromosome contacts which have been obtained from the matrix shown in **a)**. The blue regions are plectoneme free regions (PFRs). These regions are characterized from regions with RNA-seq signals **(c)** above the cutoff value. **c)** RNA-seq data for *E. coli*. Dashed black line represents the cutoff used to demarcate PFRs (light blue bars) and PARs.

However, despite multiple efforts on the experimental front, a high-resolution picture of the 3D conformational ensemble of the *E. coli* chromosome is currently elusive. This is mostly because the precedent genome-wide experimental read-out mainly provides indirect evidence of the chromosomal organization, requiring one to infer a 3D picture by guessing from the experimental data. While classical force-field based approaches have come off ages in being able to simulate large biological assemblies and complexes,^24–26^ the system size of the *E. coli* chromosome (4.6×10^6^ bp) is significantly larger than the limit that all-atom particle-based models or even systematic coarse-grained model such as Martini^27–31^ can offer. More importantly, sampling a detailed model of such magnitude via Newtonian mechanics is not tenable for a long time-scale as demanded by a heterogeneous system. On the other spectrum, phenomenological model based on polymer physics^11^ are too simplistic to capture the biological self-organization.

In the present article, we take a data-driven route to develop a quantitative 3D model of the conformational ensemble of *E. Coli* chromosome. Being aware of possible caveats of simulating the large sizes associated with modeling such systems, here we take an integrative modeling approach. In particular we combine multiple experimentally derived data of diverse resolution within a polymer based framework. We base our current modeling effort on the hypothesis that the bacterial chromosome organization is a result of fine balance between two aforementioned factors, namely intra-chromosomal contacts (Figure 1 a) and DNA supercoiling as manifested by plectonemes (Figure 1b-c). Towards this end, in this article, we present our development initiative of computer model of bacterial chromosome that integrates experimentally derived measurements of genomic contact probabilities with that of plectonemes. In particular, we combine recently reported Hi-C data of E. Coli chromosome^17^ at 5000 bp with RNA-seq data^23^ at near base pair resolution within a polymer-based framework to derive a 500 base-pair resolution model of the same. Furthermore, the bacterial cytoplasm has also been taken into account via explicit incorporation and interaction optimization of ribosomes with DNA using experimentally reported linear densities, masses and sizes. Finally, the simulations take place within a spherocyllindrical confinement whose size is commensurate with experimentally determined cell sizes. The conformational ensemble thus decorated by ribosome-sized particles within a cytoplasmic confinement and sampled by dynamical simulations, provide a quantitative, high resolution model of the bacterial chromosome integrated with experimental data at different length scales and resolutions.

## 2 Methods

To build a quantitative model for *E. coli* chromosome, we follow an integrative modeling approach via incorporation of multiple structural experimental data available in a polymer-physics based framework. The experimental data are genome-spanning and comes at multiple resolutions. In particular, two recently reported experimental measurements form the base of the current investigation: i) Hi-C derived genomic contact probability map (at 5000 bp resolution) reported in 2018^17^ and ii) RNA-seq data reported in 2019.^23^ Figure 2 illustrates the full protocol adopted here for deriving this model. Below we summarize our modelling approach in following steps, each of which we have detailed in multiple subsections.

**Figure 2:**
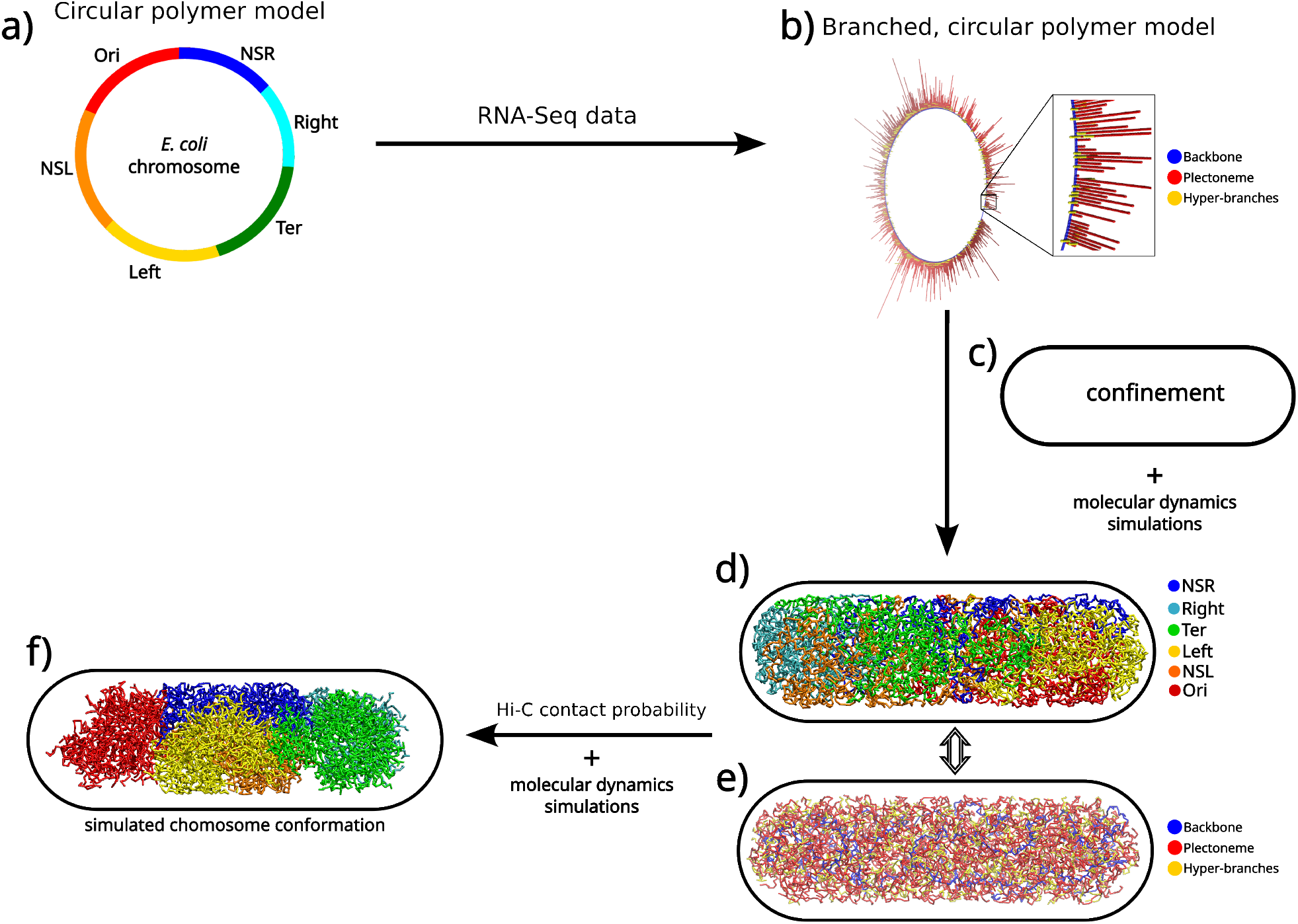
**a)** A circular polymer model with different macrodomains shown using different colours. **b)** The hyper-branched circular polymer model which was generated upon incorporation of the RNA-seq data. The chromosome beads have been classified into 3 labels: backbone (blue) which form the circular backbone, trunk (red) the plectonemes that come out as branches from the circular backbone and the hyper-branches coming out from the trunks (yellow). **c)** The chromosome is confined into a capsule shaped volume which mimics the cell wall. **d)** Chromosome conformation obtained after simulation showing the different macrodomains. **e)** Chromosome conformation obtained after simulation showing the three types of beads. **f)** The final simulated chromosome conformation.

1. A bead-in-a-spring circular polymer model forms the basic backbone of the model chromosome at 500 bp resolution.
2. We assign the labels “Plectoneme Abundant” or Plectoneme Free” to regions of the chromosome using a recently reported RNA-Seq data^23^ and the number of plectoneme abundant regions that have been experimentally observed. ^32^
3. The lengths of the plectonemes are determined such that they follow the experimental plectoneme length distribution.^33^
4. We next generate the initial configuration of the chromosome for simulations using the above mentioned experimental data and an indigenous algorithm which has been detailed in a later section.
5. Experimentally measured 5000 bp-resolution Hi-C contact probabilities^17^ are then encoded via introduction of harmonic bonds.

4. Once Hi-C has been incorporated, we model the confined cytoplasm via introducing ribosomes into the cell.
5. We parameterize the non-bonded interactions among DNA-DNA, DNA-Ribosome and Ribosome-Ribosome via iterative optimization. We also define the complete Hamiltonian for the system in this step.
6. With the architecture and the force-field ready, we perform dynamical simulations to generate a large ensemble of chromosome conformations.

### 2.1 Mapping of the bacterial chromosome topology in a circular polymer backbone

An unreplicated form of *E. Coli* chromosome forms the centre of current modelling initiative. The *E. Coli* genome has a circular topology with 4.64 × 10^6^ bp of DNA present in its unreplicated form. Here we map this chromosome in a bead-in-a-spring polymer with each bead representing 500 bp of nucleotide in a circular topology. (see Figure 2a). The choice of 500 bp as the resolution of the model lies between the resolution of experimentally derived Hi-C contact map (5000 bp) and that of RNA-seq data (∼1 bp) and serves as a judicious trade-off of between accuracy and efficiency. The 500 bp nucleotides are indexed and annotated as per the genetic sequence of wild type E. coli MG1655 and modeled as non-overlapping van der Waals particles in a polymer.

It was experimentally observed that *E. coli* cells almost always contain chromosomes which are partially replicated.^34^ Each replicated region, until the replication is completed, is termed as a replication fork. Here we redefine a parameter called the Genome equivalent (G)^34^ of a cell given by Eq-1 which is an indication of the replication status of the chromosome.

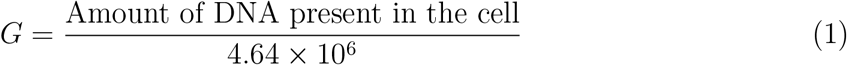

For *E. coli* in good growth conditions, multiple replication forks appear for each chromosome which results in a complicated architecture that can pose certain modeling challenges. But for moderate or slow growth conditions, a single or no replication fork was observed. ^34^ Therefore for ease of initial modeling and validation, we considered an *E. coli* cell in moderate growth condition (30 ^◦^C in M9 Minimal Media) containing a single chromosome with no replication fork (*G* = 1.0). Thus, the number of beads present in the polymer modeling the unreplicated chromosome is 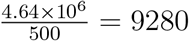. Since cell dimensions of *E. coli* also depend on the growth conditions, we modeled chromosome to be confined within a sphero-cylinder with a diameter equivalent to 0.82*µm* and total length (including end-caps) of 3.05 *µm* which are the experimentally observed average cell dimensions. As would be presented near the end of the article, as an extension of the core model, we have also developed a model for a partially replicated chromosome (*G* = 1.6) via introducing a replication fork.

### 2.2 Annotation of chromosome regions as either plectoneme abundant or plectoneme free

We have used the RNA-seq data on *E. coli* ^23^ to ascertain which regions have plectonemes. Plectonemes are formed when the DNA is supercoiled (Figure 1). It is a way by which the bacteria compacts DNA. However transcription machinery cannot read supercoiled DNA.^35^ Hence regions which contain genes that need to be transcribed are not supercoiled.^36^ This gives rise to sections of the DNA with high plectoneme density (referred as Plectoneme Abundant Regions (PARs)) connected via regions with no plectonemes (referred as Plectoneme Free Regions (PFRs)) (Figure 1). RNA-seq data provides the information on which regions of the chromosome are being transcribed. High RNA-Seq intensities indicate regions with high transcription. Therefore regions with high RNA-seq signal intensities are not supposed to contain plectonemes.

However we chose a model resolution of 500 bp per bead to strike a balance between accuracy versus computational expense while the RNA-seq data has a resolution in base pairs. Therefore, to incorporate the RNA-seq data at the chosen resolution, we ‘coarsegrain’ the RNA-seq data such that it represents the mean extent of transcription in each 500 bp region of the chromosome as described below.

Let the RNA-Seq signals be represented by {*s*_*i*_(*g*_*i*_)} appearing at genomic locations {*g*_*i*_} where *i* = 1 to *N*_*RNA*−*seq*_ where *N*_*RNA*−*seq*_ is the number of RNA-seq signals. We club together signals appearing in intervals of 500 bp and calculate their means to obtain the coarse grained signal (Eq-2).

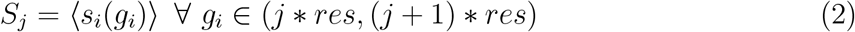

where *res* is the resolution of the model in base pairs, *j* = 1 to*N*_*beads*_ and 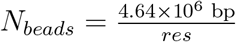 For our model *res* = 500 bp and *N*_*beads*_ = 9280.

Subsequently the RNA-seq signals are normalized within a scale of 0 to 1. To determine the threshold signal strength cutoff which sets apart plectoneme abundant regions (PARs) from plectoneme free regions (PFRs), we utilized the reports by Sinden et. al^32^ that the number of plectonemic regions per genome equivalent of DNA is 43±10. Towards this end, first we labeled all regions as plectoneme abundant (PAR). Then we labelled regions containing rRNA operons as plectoneme free as they are one of the most transcribed regions of the chromosome. Then we iteratively tuned the cutoff (= 0.33) till the number of plectonemic regions was between the experimental range^32^ (Figure S1). Once we have obtained the desired cutoff, we label the polymer beads as belonging to either PAR or PFR. To ascertain the consistency of integration of the RNA-Seq data into our model, we had also processed a previously reported RNAP-ChIP data^22^ which provides information similar to RNA-Seq. We observe a lot of similarity in the generated plectoneme abundant regions using both the data (Figure S2). However, since the RNA-Seq data has more recently been reported than the RNAP-ChIP data, we integrated the RNA-Seq data into our final model.

### 2.3 Generation of plectoneme lengths following an experimentally observed exponential distribution

The plectonemes,formed due to supercoiling, have finite lengths. It was reported by Boles et. al that the average plectoneme length is about 10 Kbp and the lengths follow an exponential distribution.^33^ Therefore once we have demarcated which regions across the circular chromosome would have plectonemes, we calculate the lengths of the plectonemes such that they follow an exponential distribution with a mean of ∼10 Kbp. We fix the minimum (*L*_*min*_) and maximum length (*L*_*max*_) a plectoneme can have to values of *L*_*min*_ = 4 Kbp and *L*_*max*_ = 75 Kbp. The minimum value has been assigned as 4Kbp as we wanted to have such a situation where even the smallest plectoneme (4 Kbp ≡ 8 beads) can also have a hyper-branch coming out of it. The choice of *L*_*max*_ on the other hand is mostly arbirtrary. Since the lengths are drawn randomly from an exponential distribution, the lengths assigned are stochastic in nature. This goes hand in hand with the stochastic binding of loop-forming proteins such as HU and MukBEF.^37,38^ Both proteins do not have any specific binding motif and bind throughout the chromosome without any specificity. Thus the loops formed by them, which are then supercoiled into plectonemes, also have lengths which are stochastic in nature.

### 2.4 Incorporation of branches as plectonemes

As discussed earlier, we have labeled regions of the chromosome as PARs and PFRs. In the polymer model, we have denoted the PFRs in light blue and the PARs in light red (Figure 3a). Since formation of branches rearranges the bonds in the polymer, we map each bead of the polymer to a genomic region for referencing to them later. We then rearrange the beads and the bonds in the circular polymer to generate a hyper-branched polymer with a circular backbone. We do so by first dividing the PARs into plectonemes whose lengths follow an exponential distribution as discussed in the previous section (Figure 3b). Each set of plectonemic beads is then converted into a branch (red) (Figure 3c) such that one bead still remains in the backbone (red beads in-line with the blue beads in Figure 3c), while the rest protrude out from the backbone. The bead remaining in the backbone is then re-labeled as belonging to the backbone (conversion of the red colored beads in question into blue beads in Figure 3d). Excluding the first and the last beads, a few consequent beads of a plectoneme are randomly selected (Figure 3d). Those make up the hyper-branches (yellow). Out of the selected beads, one bead remains embedded in the plectoneme (yellow beads in-line with the red beads in Figure 3e) while the rest protrude out of the plectoneme to form a hyper-branch. The bead from the hyper-branch, embedded in the plectoneme, is then be re-labeled as belonging to the plectoneme (Figure 3f). This process of producing hyper-branches is repeated a few more times so that 40% of the initial total plectoneme length has been converted to hyper-branches (Figure 3f and g). The choice of how much of the branch is going to form hyper-branches (here 40%) has been arbitrarily chosen. Once this process has been completed for all plectonemes, the initial configuration is ready (Figure 3h). The beads in backbone, branches and hyper-branches are then connected as per the lines shown in Figure 3g. The algorithm thus ensures that the total number of beads in the model remains constant while maintaining the genomic continuity and introducing sufficient hyper-branching.

**Figure 3:**
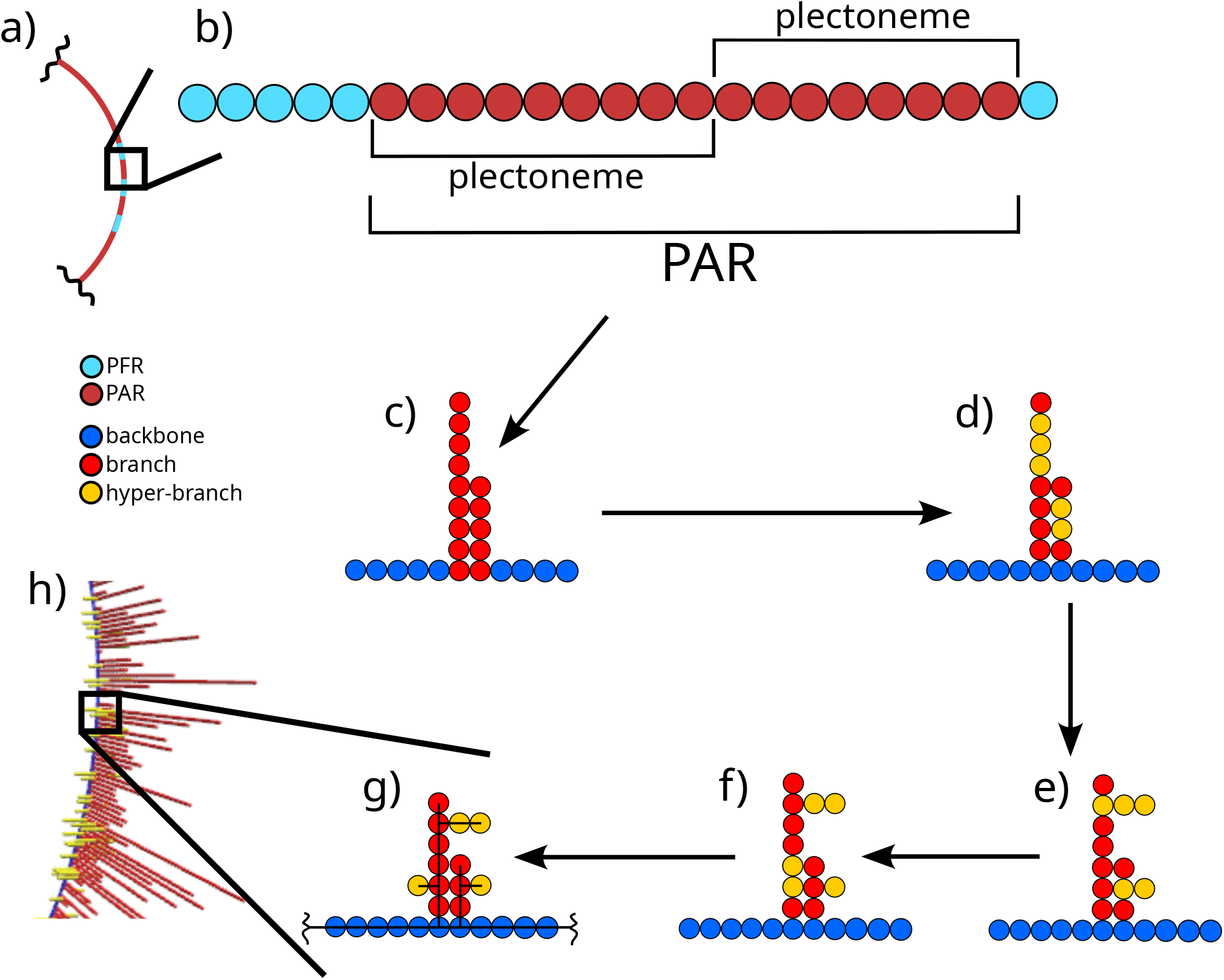
A scheme outlining the algorithm to generate the initial configuration that maintains the actual chromosome architecture determined from RNA-seq and plectoneme length distribution data. A change in color of the same bead means it has been absorbed/converted into another type of bead. **a)** A region of the circular polymer. The polymer has been annotated and mapped to the regions of the *E. coli* genome later U00096.3. ^39^ The color coding shows the classification of chromosome regions into PFR and PAR. **b)** A zoomed in part showing the beads of the polymer as PAR or PFR. The PAR has been shown to be composed of two “to-be” plectonemes. One bead from each of the branch has also been included into the circular backbone. **c)** The PAR has been divided into two plectonemes which come out as branches from a circular backbone. **d)** A few beads (in yellow) have been selected to form hyper-branches. **e)** The hyper-branches have been formed. **f)** One bead from each hyper branch has been absorbed into the branch and a few beads again are selected to form another hyper-branch for a plectoneme. **g** Another hyper-branch is formed. Then the beads are connected via harmonic bonds shown in black lines passing through the beads. **h)** Shows a section of the actual initial chromosome configuration generated which is a zoomed out image of **(g)**.

While the majority of the model centers around an unreplicated chromosome (G=1.0), we have also simulated a partially replicated chromosome by introducing a replication fork. Towards this end, we have used a similar approach as employed in our previous work^40^ by copying the architecture of the mother DNA that has been replicated and connecting it to the main chain. The points where the replicated region and the mother DNA meet are termed as the fork points. However we made sure that the fork points remains plectoneme free as replication also requires the DNA to be plectoneme free.

### 2.5 Encoding Hi-C-derived contacts as harmonic restraints

The Hi-C-derived contacts have been encoded into our polymer as harmonic distance restraints. Considering the 5 kbp resolution of the reported Hi-C contact map, we incorporated distance restraints at the 5-th bead of each 10-bead segment of the polymer irrespective of the identity of the beads. Let these beads be called “Hi-C beads”. If we label the beads of the polymer beads, as per their genomic coordinate, from 0 to *N*_*beads*_−1, where *N*_*beads*_ = 9280, the Hi-C beads are then constituted by beads 5, 10, 15 and so on. Thus there are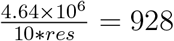 beads since *res* = 500. This is equal to the dimension of the experimental Hi-C contact probability matrix. Thus the maximum number of possible Hi-C pairs that can be encoded is ^928^*C*_2_. The conversion of Hi-C contact probability matrix into distance restraint is similar to our previous approach.^40^

We first assumed that the distances between the Hi-C beads 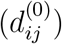 is equal to 1/*p*_*ij*_ where *p*_*ij*_ is the contact probability matrix between “Hi-C beads” *i* and *j*. To maintain these distances we incorporate harmonic springs between these beads. The force constants (*k*_*ij*_) of the springs are given by *k*_0_ exp 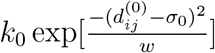 where *k*_0_ determines the maximum stiffness of the springs. *w* is a parameter that needs to be tuned to reproduce the experimental Hi-C matrix from the simulations. However, the experimental Hi-C matrix is sparse and a lot of its values are zero or very low. For those probability values, the distances are very large (greater than the cell length). Also the values of *k*_0_ exp 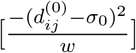 are very low for such values. Such low force constants do not affect the chromosome conformation significantly and can be safely ignored. Therefore finally we incorporate only those Hi-C contact probabilities whose respective *k*_*ij*_*/k*_0_ values are higher than 10^−3^. Here, *w* is parameter that affects the reproducibility of the simulated contact probability matrix and needs to be optimized. We found that *w* = 1.0 provides the optimum similarity in experimental and simulated contact probability matrices where only about ∼5% of the total number of possible beads pairs are connected via Hi-C bonds (Figure 4b).

**Figure 4:**
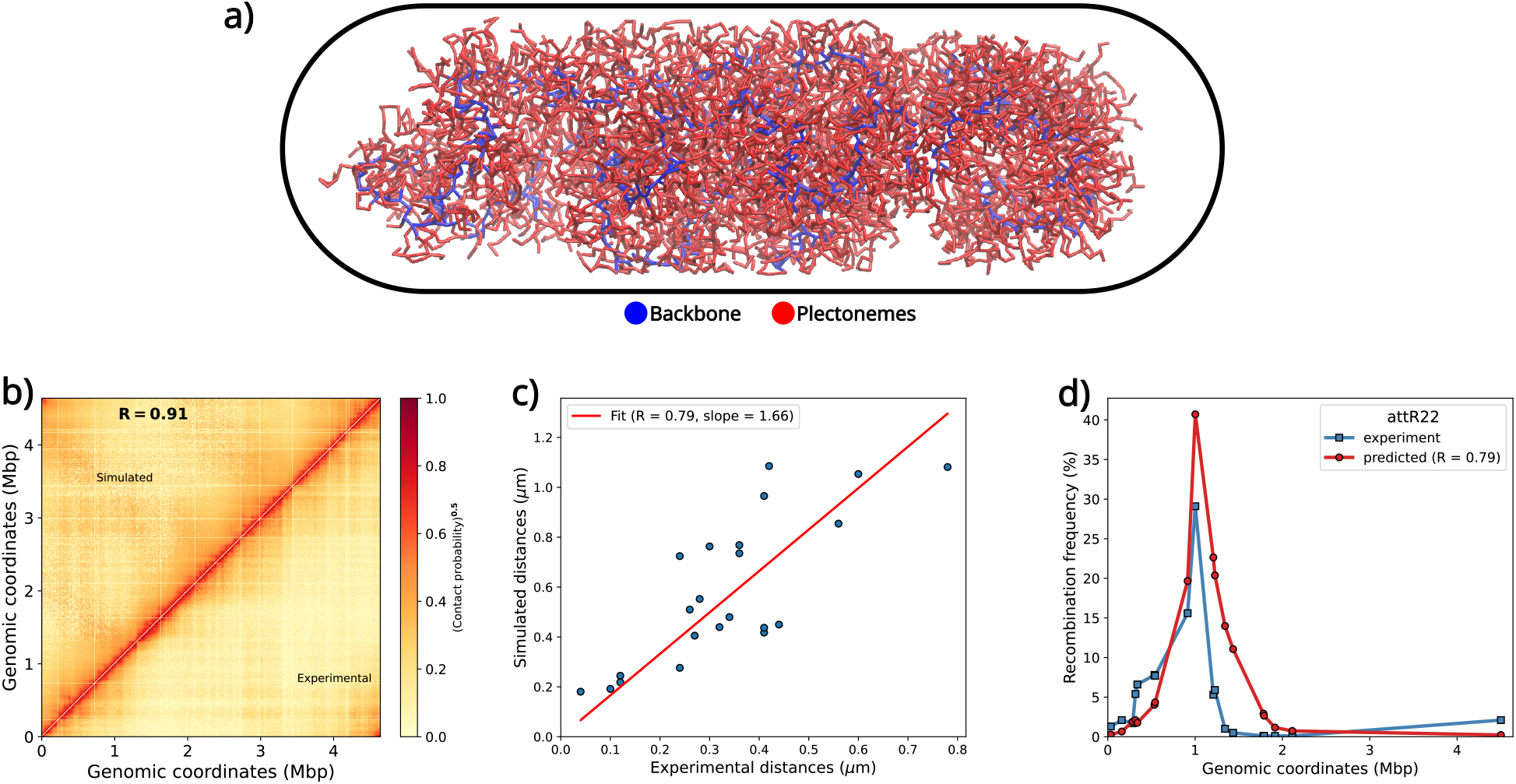
**a)** Representative chromosome conformations obtained via simulations. The different colors represent plectonemes and the backbone. **b)** The regenerated contact probability matrix. The upper diagonal depicts the matrix regenerated from simulations while the bottom diagonal shows the experimental contact probabilities for a direct comparison. The values of each contact have been enhanced by raising each value to the power 0.5. **c)** Comparison of simulated and experimental distances among a select loci pairs. The red line shows the *y* = *mx* fit. **d)** Comparison between recombination frequencies predicted from simulations (red) with the experimentally reported values(blue).

It should be noted that for architectures with replication forks, we do not include any Hi-C restraints between beads of daughter and mother DNA, consistent with our previous related investigation.^40^

### 2.6 Modelling the cytoplasm of *E. coli*

The cytoplasm of *E. coli* consists of many different species of biomolecules. However, one of the most abundant and volumetrically large species are the ribosomes. In the present work, we model the cytoplasm by introducing adequate number of ribosomes commensurate with an *E. coli* cell, growing in moderate growth conditions. We incorporate both monomeric ribosomes in the form of 30S and 50S, as well as polymeric ribosomes (referred here as polysomes) with each subunit of 13-mer polysome representing 70S a ribosomes. The total number of ribosomes estimated to be present in the present model, is ∼21000^41^ with a polysome-monomer ratio of 80:20.^11^ Therefore, there would be 2102 30S subunits, 2102 50S subunits and 1292 polysomes. ^42^

We have modeled a single ribosomes subunit as a sphere. The size of the sphere depends on its identity as 30S, 50S or 70S. The sizes and masses assigned to the spheres are as per experimentally reported numbers (Table S1). There are also bonds present between adjacent 70S beads of the polysomes. We shall discuss the interactions in details in later sections. However, to avoid digression, we postpone any discussion pertaining to ribosomal particles for a future article.

### 2.7 Description of the Interaction Hamiltonian of the model

The Hamiltonian of the system is given by Eq-3.

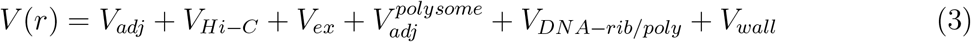

where the first two terms are exclusively for the chromosome. *V*_*ex*_ is a generic volume exclusion term which ensures the systems faces a penalty if particles start overlapping. 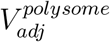 is the potential between adjacent 70S subunits that constitute a polysome. *V*_*DNA*−*rib/poly*_ represents the interactions among chromosome and ribosomes. Finally, *V*_*wall*_ enforces the confinement due to cell wall. We shall discuss each potential term in detail in this section to provide the rationale behind each term.

We have assumed that DNA-DNA and ribosome-ribosome interactions are purely repulsive in nature defined via Eq-4. Such interactions are simply short ranged, excluded volume interactions in nature.

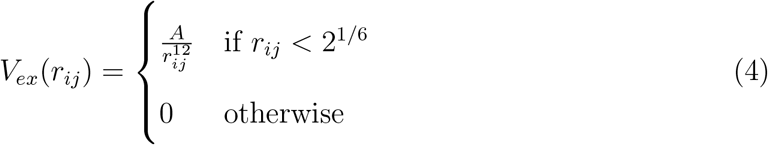

The value of *A* for different particle pairs (DNA-DNA, polysome-polysome, polysome-ribosome and ribosome-ribosome) have been reported in Table-S2.

However, we have not assumed that DNA and ribosome interactions are also volume exclusion interactions, on the contrary to a previous model.^11^ We have the potential defined in Eq-5.

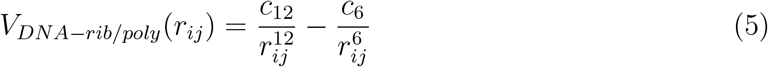

where *c*_6_ and *c*_12_ have been tuned to match the simulated and experimental linear density profiles of DNA and ribosomes and have been reported in Table-S2.

The adjacent beads of the polymer, that models the chromosome, are bound via harmonic springs defined by Eq-6.

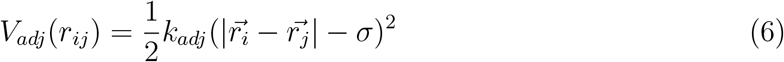

where *V*_*adj*_(*r*_*ij*_) is the potential that binds adjacent beads of the polymer and *k*_*adj*_ is the force constant of the springs connecting adjacent beads of the polymer. 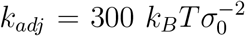 has been used.

A similar type of springs, however much stiffer, binds the adjacent beads of the polysome. It is given by Eq-7.

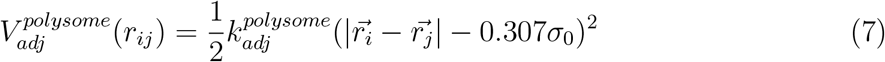

where 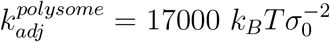.

The “Hi-C beads” defined in the previous section are also bound via harmonic springs which is given by Eq-8.

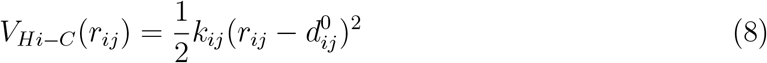

where *V*_*Hi*−*C*_(*r*_*ij*_) is the potential that maintains proper distances among the Hi-C beads obtained from experimental contact probability matrix and 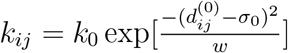.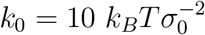 has been used for this study.

Finally to mimic the cell wall, we have also incorporated a confinement potential that acts inwards whenever a particle tries to cross the cell wall. The potential is given by Eq-9.

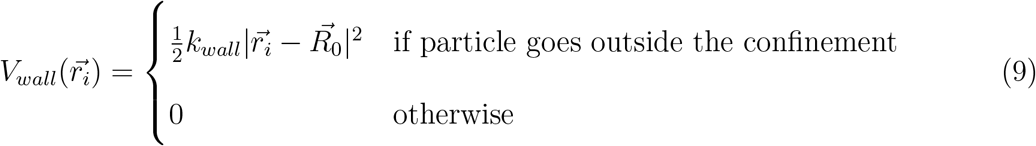

where 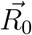 has been set as the center of the confinement and 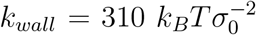 has been used.

### 2.8 Simulation protocol

We generate 32 independent initial configurations of chromosome following the architecture that we extracted form chromosome and plectoneme related structural experimental data.^23,33^ It is to be noted that we have not incorporated the contact probability data into the model yet. Then we introduce the required number of ribosomes and polysome.^41^ These serve as the initial configurations.

Upon successful generation of an initial configuration of the chromosome, we turn on the interactions that maintain the distances among different regions of the chromosome obtained from Hi-C contact probability matrix. Then we run an energy minimization with a force tolerance criteria of 10 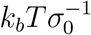. Next we run molecular dynamics simulations using the Velocity-Verlet integrator for 3×10^6^ steps with a time-step of 0.002 *τ*_*MD*_ at *k*_*B*_*T* = 1 where 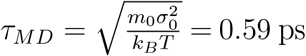. All the simulations have been performed using a modified version of GROMACS 5.0.6^43,44^ where we have implemented the spherocylindrical confinement (https://github.com/JMLab-tifrh/gromacs_spherocylinder). For all analysis we use the last 1000 frames of the generated trajectories.

## 3 Results

Here, using a series of protocol (see Figure 1 and Figure 2), we develop a model of *E. coli* chromosome by integrating two key experimentally measured genome-wide data within a polymer-based architecture: Hi-C derived intra-genome contact probability map^17^ measured at a 5000 bp resolution and RNA-seq data^23^ measured at a 1 bp resolution. RNA-seq data, quantitatively informs one with the locations of high and low transcribed regions across the chromosome. In the current investigation we use the idea that transcription machinery cannot read supercoiled DNA. ^35^ (see Figure 1b and *Methods*). With the knowledge of the number of plectonemic regions that should be maintained^32^ and the length distribution of the plectoenemes^45^ as additional constraints, we generate a hyper-branched ring polymer topology of chromosome model with optimal assignment of PAR and PFR along the chromosome (see *Methods* for details). On the other hand, the data derived from chromosome conformation capture experiment forms the other key aspect of the model. In particular, we introduce recently reported chromosome contact probability matrix as derived from Hi-C data pertaining to *E. Coli* chromosome,^17^ at a resolution of 5000 bp. Towards this end, we employ a protocol recently implemented by our group^40^ where the experimentally obtained contact probabilities are incorporated as harmonic restraints into the hyper-branched topology of chromosome. The joint implementation of these two important key genomic data at two different resolution, together with appropriate confinement, equivalent to actual cell dimensions, and other related experimental data^11,12,34,46^ (see Figure 2 for protocols) produces a 500 bp-resolution chromosome conformation.

As it follows in rest of this section:

- we first present validation of the constructed chromosome model against experimental data.
- Next the chromosome model is characterized for its hierarchical organization.
- Finally we demonstrate a set of applications of the model for prediction and interpretation.

### 3.1 Validation of the chromosome model against multiple experimental data

A representative conformation of the chromosome obtained via the simulations have been shown in Figure 4a. The blue color represents the backbone out of which plectonemes (red) originate. For *C. cresentus*, it was observed via experimental contact probability maps and computational modeling that the chromosome adopts a bottle-brush like structure having a central core with the plectonemes radially protruding from it.^47,48^ A recent experiment^49^ hints towards such an architecture for *E. coli* also.^50^ Incidentally, the chromosome conformation, obtained using the current protocol, looks like a typical bottle brush-like structure, resembling the emanation of branches as a direct outcome of plectonemic features. However, before we begin to analyze the structural aspects of the plectonemes, we first verify the quality of the present model against a set of experimental data. First we ascertain whether the chromosome contacts are properly maintained in the model derived via integration of Hi-C and RNA-seq data. As we encoded only a small amount of the contact probabilities as harmonic restraints, we first reconstruct the full contact probability matrix at 5000 bp resolution (same resolution as reported in experiment^17^) via averaging over the ensemble of simulated chromosome conformations.

We do so by calculating the distance matrix between any two polymer beads at an interval of 10 beads (5000 bp) from each frame, denoted by *D*_*i*_. For each distance matrix, we converted them to a contact probability matrix using 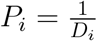. The final contact probability matrix is an average of all probability matrices obtained from each frame and each seed, i.e 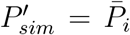. Next we filter 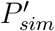 as the experimental matrix contains zeros, most of which tend to zero in our simulated contact probability matrix, but are not exactly zero (Figure S3a). That is because it would require ∞ distance between two regions to get zero contact probability between two regions, as we have assume that 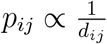. We do so as per Eq-10.

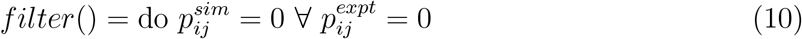

The filtering simply locates which elements in the experimental matrix are zero and replaces the respective values in the simulated matrix with zeros. The final filtered matrix is given by 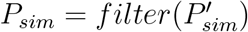.

From Figure 4b, we can see that simulated model have been able to reproduce the complete experimental contact probability matrix to a high degree of similarity as per the Pearson correlation coefficient observed between the simulated (*P*_*sim*_) and the experimental (*P*_*expt*_) matrices. For a more quantitative comparison, we plot the difference (*P*_*sim*_ − *P*_*expt*_) matrix in Figure S3b. We find that most values lie near zero. To conclusively prove that we plot a distribution of the absolute values of the difference matrix, while neglecting the zeros (Figure S3c). The distribution shows the most values of the difference matrix lie between 0.051 and 0.081 which are really low in magnitude. The low MSE subtly augments the similarity in the reproduced simulated and the experimental matrix and supports our claim that the similarity is not only qualitative but also quantitative in nature, i.e almost exact reproduction.

For a more rigorous assignment, we also compared already reported distances obtained via fluorescence microscopy^18^ (which have not been used for developing the present model) and the distances obtained via the current model (Figure 4c). We plotted the experimentally observed distances along the x-axis while the distances obtained via simulations along the y-axis. We report a Pearson correlation coefficient between distances obtained via experiments and simulations (=0.79). Considering the fact that our model has not been introduced to this data at all during modelling, we consider the correlation to be quite encouraging. We also found the slope of the fit to be *m* = 1.66, implying that though the trend in the distances among the loci remains similar to the trend observed in the experiments, the distances among all loci pairs are higher in the simulations. This can be due to differences in cell lengths and chromosome contents.

The simulated recombination frequency data for attR22 are compared with the experimentally obtained values.^20^ To be able to predict the recombination values for the attR22 probe, we first calculate the spatial distances among all probe pairs. We plot the spatial distances along the x-axis and plotted their corresponding observed recombination frequency along the y-axis (Figure S4). We then fitted them using an exponential function (red line in Figure S4). For the attR22 probe pairs, we first calculate their genomic and spatial distances. From the obtained spatial distances we predict their probable recombination frequency using the parameters obtained via the exponential fit in Figure S4. We plot the predicted recombination frequency vs the genomic distances in Figure 4d in red. In blue we show the recombination frequencies observed experimentally and we find a good qualitative agreement between experiment and predictions (R=0.79).

### 3.2 Characterization of the chromosome model

#### 3.2.1 The chromosome model captures macrodomain spatial segregation properly

One of the hallmark features of the *E. coli* chromosome is the segregation of the conformation into four non-overlapping macrodomains (referred as *‘Ori’, ‘Ter’, ‘Left’, ‘Right’*) and two non-structured regions (*‘NSR’ and ‘NSL’*).^20^ Figure 5a shows the chromosome conformation where the genomic coordinates of macrodomains and non-structured regions have been annotated by different colours. We can see that no two colors mix with each other. Each occupies a specific region of space, mixing slightly only at the interface. This is due to the separation of different macrodomains and non-structured regions into their own spatial regions as predicted by recombination assay experiments.^20^

**Figure 5:**
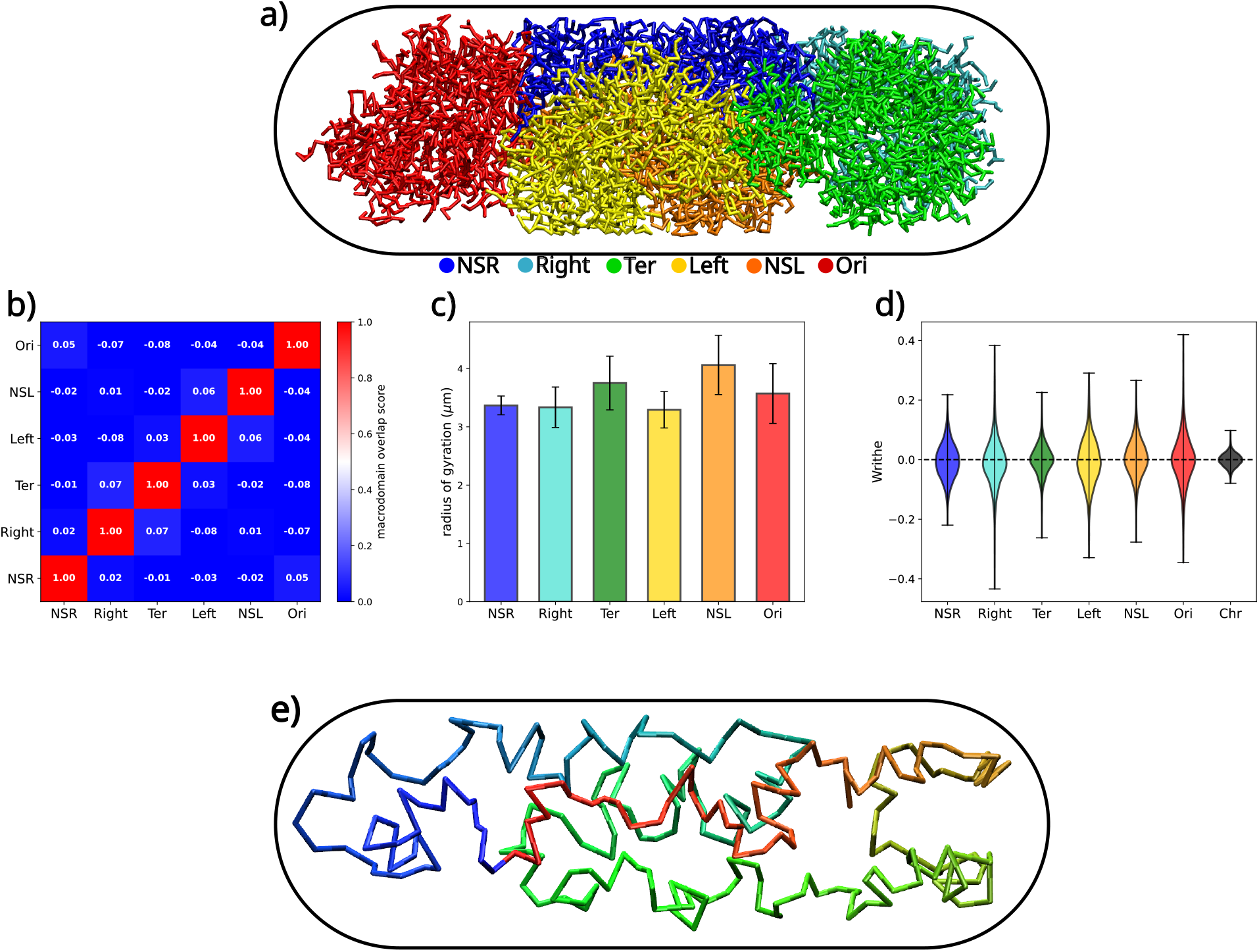
**a)** Representative snapshot for the chromosome showcasing the different macrodomains and non-structured regions. **b)** Macrodomain density correlation matrix. Blue means no correlation or no mixing between those macrodomains. Red means complete mixing. **c)** Radius of gyration of different macrodomains and non-structured regions. **d)** Writhe values of all the macrodomains and non-structured regions. **e)** A representation showing the helical organization of the chromosome.

To quantify the segregation of macrodomains observed in Figure 5a, we have calculated the 3D density correlations of macrodomains inside the cell. First we divided the volume of the cell into small cubes, called voxels. Considering only one macrodomain at a time, we calculated the number density of the macrodomain in each voxel. If two macrodomains mutually overlap, then these would have some voxels where both have non-zero densities. If they do not mix at all, then they would not have any voxel where both would obtain nonzero density simultaneously. This property has been used to calculate macrodomain overlap scores in Figure 5b. The overlap score is simply the Pearson correlation coefficient between density tensors obtained for two macrodomain pairs. For non-overlapping macrodomains, the Pearson correlation coefficient would be close to zero, while for fully mixed macrodomains it would be one. As a proof of concept we have also reported the self-overlap score for each macrodomain which can be seen from the diagonal which has a value of 1. Overlap scores for all macrodomain pairs are near to zero as can be seen from the blue colour of the non-diagonal regions in Figure 5b.

Next we calculate the radius of gyration (*R*_*g*_) of each macrodomain for a quantitative measure of size. Hence from the calculated radius of gyration, we can obtain an estimate of the size of each macrodomain and the non-structured regions. From Figure 5c, it can be seen that NSL and Ter have the highest values of *R*_*g*_ followed closely by Ori. NSR, Right and Left have very similar sizes. The values have a similar trend is very similar to the trend of the *R*_*g*_s obtained by Hacker et. al for their plectonemic model with oriC at mid cell. ^51^ However we do not have a fixed disposition of oriC in our model and it actually has a distribution where it is located either at mid-cell or the cell pole (Figure S7).

Finally we calculate the writhe of the whole chromosome and also each macrodomain to investigate the overall chiral disposition of the chromosome. We find that the writhe of each macrodomain is centred around zero with a standard deviation much greater than the mean (Figure 5c). As per previous experimental investigations into the chirality of the full chromosome,^52^ it was seen that such a trend, i.e. a near zero mean and a high standard deviation, signifies a helical organization of the chromosome with no net chirality. As we find similar trends in the writhe for all macrodomains, we think that each macrodomain is also helically arranged inside the bacterial cell, however with no net chiral disposition. The same is shown via Figure 5e.

#### 3.2.2 Plectoneme free regions remain closer to the periphery of the bacterial cell wall

The resolution of the present model provides us an opportunity to statistically characterize the location of plectoneme across the bacterial genome. First, to verify whether our attempts to enforce proper plectoneme lengths in our model that follow the previously reported exponential distributions with a mean of 10 Kbp^33^, we plot the individual plectoneme length distributions obtained from each of the 32 independent realizations in Figure 6a. The black line is the reference line which has been constructed numerically. We see that all the non-black lines closely follow the black line with a collective mean of 9.22 Kbp, in accordance with experimental report.^33^

**Figure 6:**
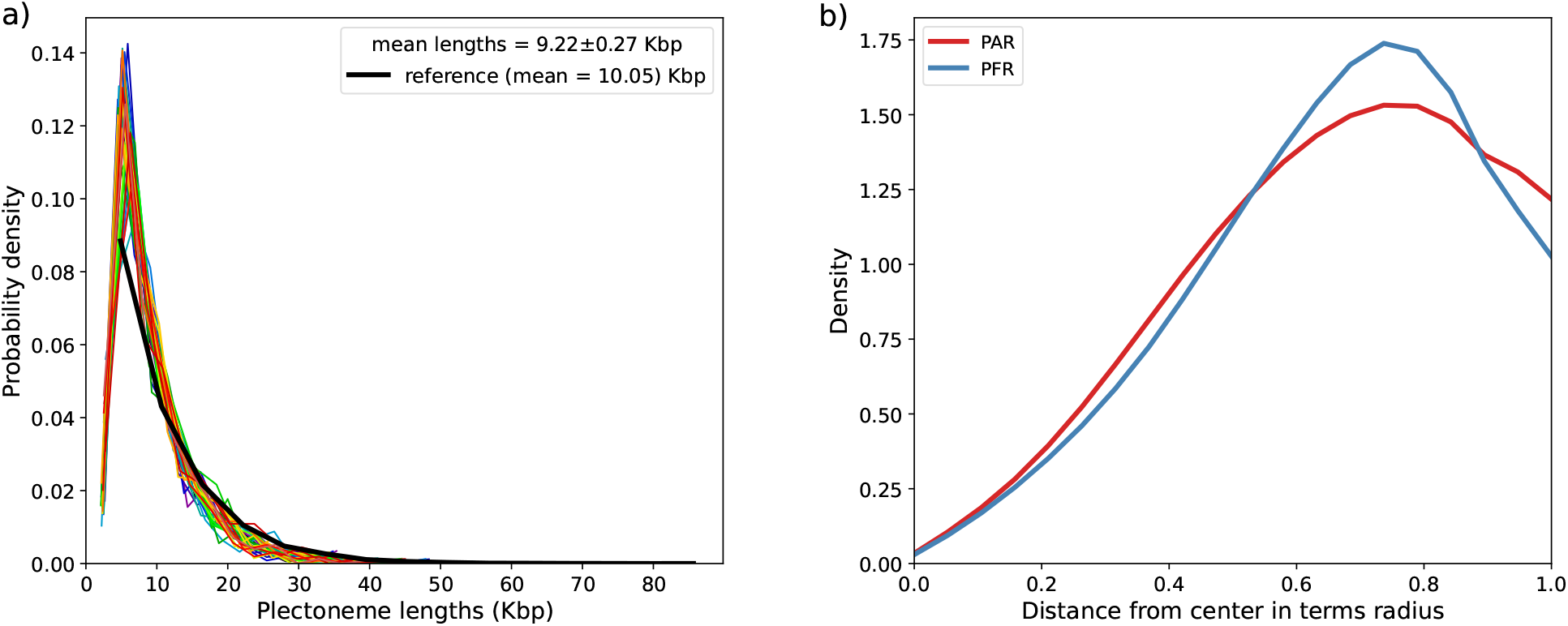
**a)** Length distribution of plectonemes in the model. **b)** radial disposition of PARs (red) and PFRs (blue).

Upon passing multiple validation via comparison of existing experimental data, we investigate our model-derived chromosome conformations to characterise the plectonemes and its role in genome organization. In particular, we first explore the difference in organization between plectoneme free (PFR) and plectoneme abundunt (PAR) regions. In Figure 6b we show the radial disposition of PFRs (blue) and PARs (red). The x-axis represents distance from the center of the bacterial cell along the radial direction (i.e. along short axes of the spherocylindrical cell). While both PARs and PFRs show a tendency to accumulate near the periphery of the bacterial cell, PFRs have relatively higher density near the cell-wall than PARs. The increased propensity of PFRs near the periphery is biologically relevant for survival. PFRs are regions undergoing active transcription. Nutrients entering the cell would thus be readily accessible to the PFRs if they are near the cell wall. Therefore the PFRs localize themselves in the vicinity of the cell wall which the model captures neatly.

### 3.3 Predictive Features of the model

#### 3.3.1 Prediction of a contact probability matrix higher than the experimental resolution

Using our protocol, we obtained a model that has a resolution of 500 bp which gives us an opportunity to predict a possible intra-genome contact map at a resolution higher than the current reported experimental resolution (5000 bp). Hence we calculated the contact probability matrix from our model at 500 bp (Figure 7a) for an unreplicated chromosome. From Figure 7a inset region, we can see the signatures of “microdomains” which appear near the diagonal. These patterns are formed via the following interactions: i) plectonemic architecture forces close-by regions to be densely packed resulting in an enhanced contact probability. ii) “Hi-C “ beads, when come close to each other also drag their neighbours. These neighbours also have enhanced contact probabilities. The rest of the matrix, away from the diagonal, lacks any feature.

**Figure 7:**
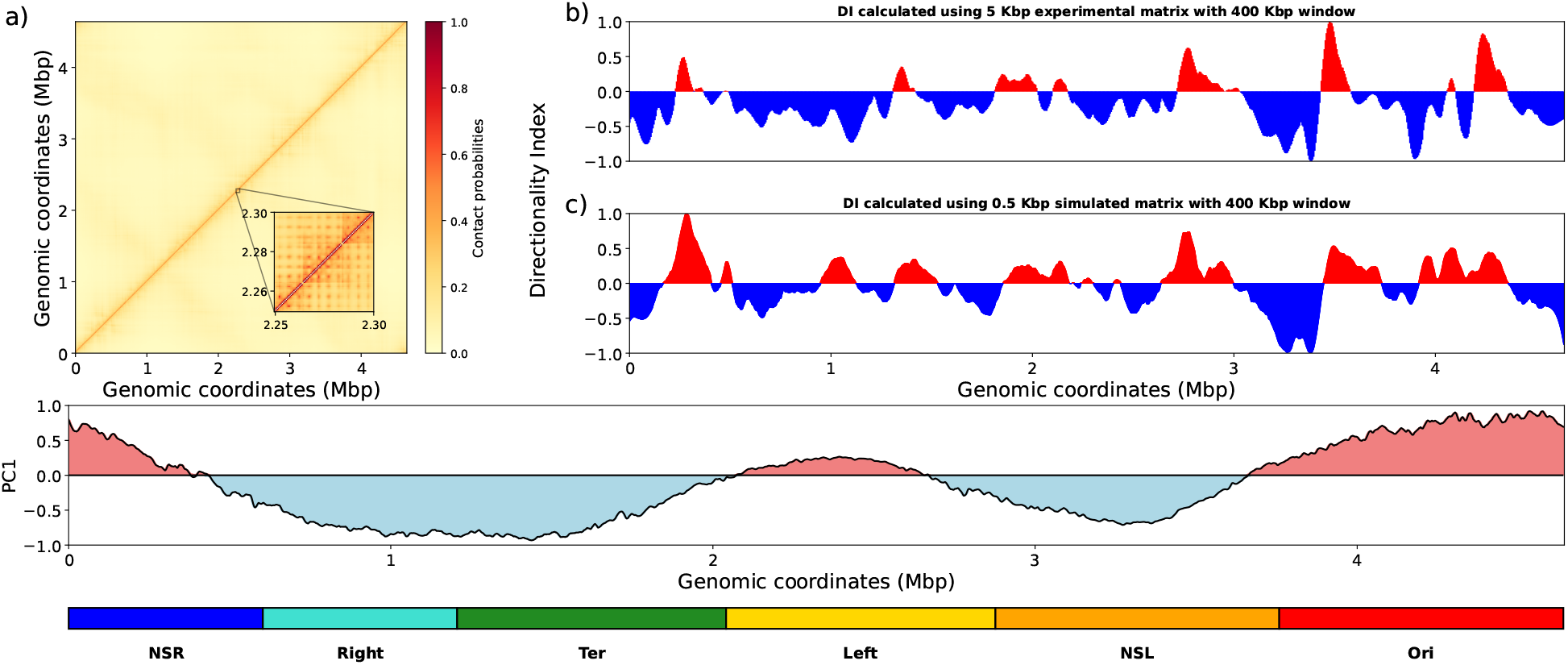
**a)** The contact probability matrix at 500 bp resolution. **b)** DI values obtained from experimental contact probability matrix at 5000 bp with a window of 400 Kbp. The DI values have been scaled between -1 and 1. Red regions indicate positive values while blue regions indicate negative values. **c)** DI values obtained from simulated contact probability matrix at 500 bp with a window of 400 Kbp. The DI values have been scaled between -1 and 1. Red regions indicate positive values while blue regions indicate negative values. **d)** Plot of the first principal component (PC1) values obtained from PCA of the simulated contact map at 500 bp vs. genomic position. Pink indicates positive values while cyan indicates negative values of PC1.

We next assess the high resolution matrix obtained from simulations. For that we calculate Directionality Index (DI) at a window of 400 Kbp using the experimental contact probability matrix at 5000 bp (Figure 7b) and the simulated contact probability matrix at 500 bp (Figure 7c). Both the DIs show a high degree of similarity. Overall both the DIs have a Pearson correlation coefficient of 0.69 which indicates good agreement between them. Therefore we claim that the contact probability matrix obtained using simulations, having a resolution of 500 bp, is fairly accurate.

Upon validation, we perform a Principal Component Analysis (PCA) on the 500 bp contact matrix and plot it against the genomic coordinates (Figure 7d). Upon overlaying the macrodomain regions, we observe that the PC1 values of adjacent macrodomains do not tend to have the same sign, except for Right and Ter. Since PC1 values in a same band means that the region tends to interact with itself rather than its neighbours^15^ and the fact that PC1 values from adjacent macrodomains have opposite signs, we conclude that the PCA decomposition of the matrix at 500 bp captures the phenomenon of macrodomain segregation which the experimental matrix at 5000 bp could not capture properly (Figure S5).

To investigate why Right and Ter did not separate out as two macrodomains, we calculate the amount of PFR present in each of the four macrodomains and non-structured regions^20^(Figure S6). A higher amount of PFR signifies a higher number of transcribing regions. Our analysis finds that Ori has the highest PFR content which means that Ori macrodomains have more number of transcribing genes than that of Ter, which has a much lower PFR content than Ori, with NSL having the least PFR content. We think that since Right and Ter are two adjacent macrodomains and also have very high plectoneme densities, it renders the simulated matrix at 500 bp unable to resolve them as different macrodomains via PCA (Figure 7d).

We also think that Ori having the highest number of PFRs hints at a functional relevance of Ori macrodomain as it might contain genes that are very basic for the survival of the organism. It can also be the reason why it gets replicated first. However these need to be experimentally verified and here these represent our opinions on the importance and possible function of Ori.

#### 3.2.2 Extending the model towards a replicating chromosome

While the current model is quite efficient in capturing the in-vivo organization of the chromosome in agreement with multiple experimental observations very accurately, the Hi-C measurements also contain contributions from cells which are replicating their chromosomes. It was seen from a previous study by Bremer and Hans, ^34^ the average amount of DNA inside an *E. coli* cell growing in moderate growth conditions was found to 1.6 genome equivalent, 1.6 times that of an unreplicated chromosome. This indicates that, in an ensemble, most cells would have undergone chromosome replication. Therefore, as an extension of the current modelling protocol we wanted to simulate a chromosome undergoing replication.

Thus we extend our model by generating polymer architectures, as described in *Methods*, to represent a chromosome with a replication fork as can be seen from Figure 8a.

**Figure 8:**
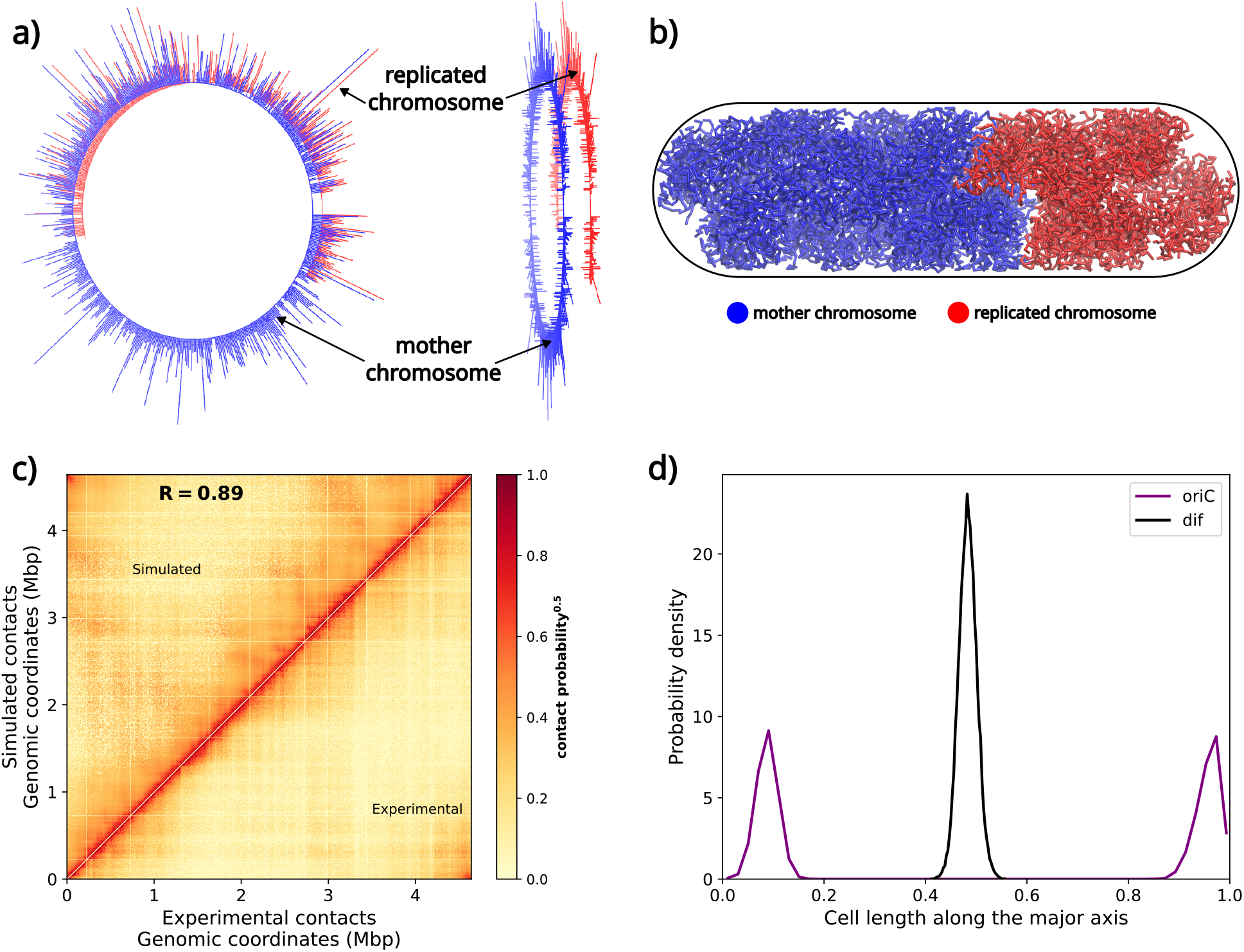
**a)** Positions of oriC (purple) and dif (black) along the long axis of the cell for an unreplicated chromosome. **b)** Representative snapshot of an unreplicated chromosome depicting the positions of oriC and dif. **c)** Positions of oriCs (purple) and dif (black) along the long axis of the cell for a partially replicated chromosome. **d)** Representative snapshot of the partially replicated chromosome depicting the positions of oriCs and dif.

The same simulation protocols as described in *Methods* have been used to simulate a replicating chromosome inside a cell. A representative conformation has been shown in Figure 8b. Using the generated ensemble of conformations, we recalculated the Hi-C contact probability matrix and compared it with the experimentally obtained matrix (Figure 8c). We observe that simulated matrix is very similar to the experimental one which signifies that Hi-C contacts encoded are properly maintained.

Next we calculated the location of oriC and dif along the long axis of the cell (Figure 8d). We observe that for our model, the two oriCs remain localised at cell quarters, consistent with previous studies.^21,40,53^ Moreover we also observe that the mother and the daughter chromosomes are properly segregated into different cell halves along the long axis (Figure S8). Such an orientation helps chromosome segregation during cell division which we obtain from our model without any external imposition.

We then calculate the number of PFRs per macrodomain for the replicating chromosome (Figure S9). We observe an increased amounts of PFRs in NSR, Right and Ori regions. We attribute this increase to two main factors: i) the replication process duplicates the mother cell’s hyper-branch architecture in the daughter chromosome and ii) replication initiates at OriC and proceeds bidirectionally with equal speeds. We also calculate the number of plectonemic regions that are present in the replicated architecture which we obtain as 69. Experiments report that for bacterial cell containing 2.8 genome equivalents of DNA (G=2.8) have 120 plectonemic regions on an average.^32^ Assuming linear dependency of plectonemic regions with the amount of DNA contained in-vivo (G value), for G=1.6, the number of plectonemic regions come out to be 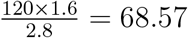. Therefore for our replicating chromosome model, the number of plectonemic regions come out to be very accurate. These prove that the protocol, along with proper polymer architecture, can generate high resolution models for the *E. coli* chromosome inside the cell for both non-replicating and a replicating chromosome with a high degree of accuracy.

#### 3.3.3 Prediction of individual role of plectoneme and chromosome contacts in organization

The results discussed above show that using multiple experimental data, we were able to successfully generate a high resolution model for the *E. coli* chromosome. However, the individual role of chromosome contacts and supercoiling in proper organization are still not clearly understood. In this respect, the modular structure of the present model provides us an opportunity to design controls that would highlight the individual roles of different features. Accordingly, we compared the WT *E. coli* chromosome conformation in its unreplicated form (Figure 9a) to three control scenarios that we computationally simulate : i) the chromosome has no Hi-C contacts but has plectoneme (Figure 9b). ii) the chromosome has Hi-C contacts encoded but has no plectoneme (Figure 9c). iii) the chromosome has neither Hi-C contacts nor plectonemes, i.e. a simple self-avoiding, circular random walk polymer (Figure 9d).

**Figure 9:**
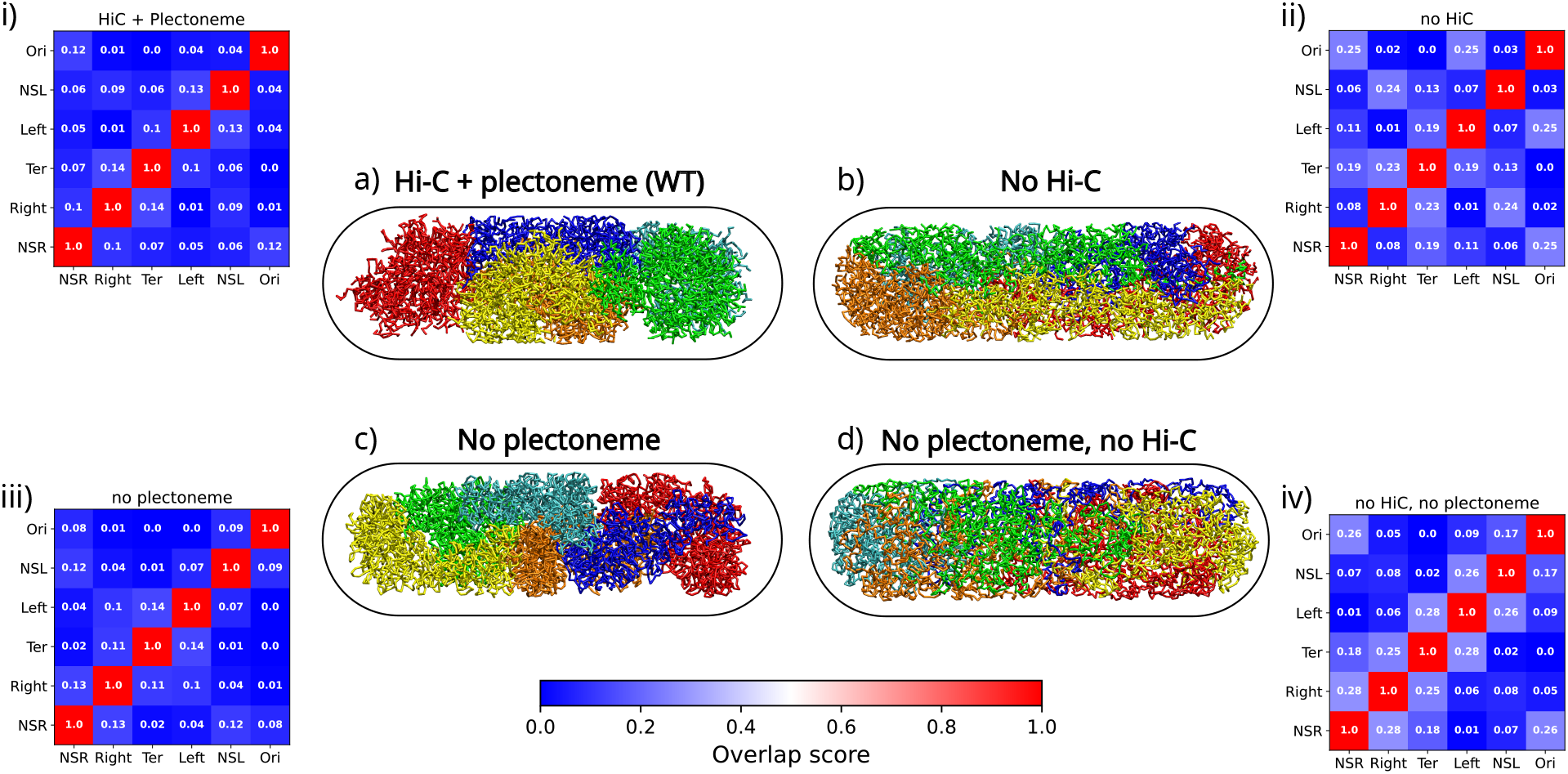
Representative snapshots for: **a)** With Hi-C and plectonemes (WT) **b)** No Hi-C, only plectonemes. **c)** With Hi-C, but no plectoneme. **d)** No Hi-C, np plectoneme. Chromosome overlap scores for: **i)** with Hi-C, with plectoneme. **ii)** no Hi-C, with plectoneme. **iii)** with Hi-C, no plectoneme. **iv)** no Hi-C, no plectoneme.

Upon comparison of the chromosome conformations in Figure 9a and Figure 9b, we observe that the macrodomains are more spread out and less compact. This is nicely captured in Figure S10 which shows that upon removal of Hi-C, all macrodomains increased in their respective sizes with respect to the WT scenario **(a)**. The macrodomain overlap scores shown in Figure 9i and Figure 9ii also indicate a relative loss in macrodomain segregation propensity. Therefore, we conclude that chromosome contacts are very important for proper compaction and organization of the chromosome and that they are one of the most important factors that maintain proper segregation of macrodomains.

Next we investigate the effect of not having plectonemes on the chromosome organization. For that we perform simulations where the chromosome is a simple ring polymer, i.e. it does not have any branches or hyperbranches that represented supercoiling. However we still retain chromosome contact information incorporated into the model. From the representative chromosome conformation (Figure 9c), we can observe that macrodomains still appear to be properly segregated and compact. However upon calculation of individual macrodomain sizes (Figure S10), we observe that the sizes of all macrodomains have slightly decreased upon removal of plectonemes. Also the macrodomain overlap scores (Figure 9iii) suggest minimal disruption in their separation propensities. This raises an interesting question on the role of plectonemes in chromosome organization. Do they have any significant role on the chromosome organization or do they function only as gene transcription regulating elements for the bacteria?

To bring forth the importance of plectonemes, we performed a set of simulations where we did neither have plectonemes nor have encoded Hi-C contact probabilities. From the representative chromosome conformation (Figure 9d), it can be seen that the chromosome organization as well as macrodomain segregation have been greatly effected (Figure 9iv). However in the presence of plectonemes but no chromosome contact information (scenario **b**), the macrodomain segregation was still present to some extent. The highly enhanced sizes of the macrodomain for scenario **d** (Figure S10) also showcase the disruption in chromosome organization which was not present for scenario **b**. Thus we conclude that plectonemes do function as chromosome condensation agents, though they form as a result of the bacteria’s efforts to regulate gene transcription. However their role in macrodomain segregation is not as prominent as that of chromosome contacts and that they enhance the effects that chromosome contact formation has on its in-vivo organization.

## 4 Conclusion

In conclusion, we have put forward a proposal for integrating multiple experimentally derived data (Hi-C contact probability and RNA-Seq data) as an input within the framework of hyper-branched circular polymer model to generate a high resolution model for the *E. coli* chromosome. The data-informed model could predict multiple, independent experimental observations and is rigorously validated against data not used during modeling purpose. The model provides a statistically robust picture of the spontaneous organization of chromosome into a set of spatially distinct macrodomains. Next we investigated the orientation and arrangement of the chromosome inside the cell. Calculation of it’s writhe showed that the chromosome is arranged in a helical manner but with no net chirality. We then use our model to investigate the arrangement of plectonemes inside the small cell volume. We observed that both plectoneme free regions (PFRs) and plectoneme abundant regions (PARs) would not like to populate the centre of the cell. However plectoneme free regions have a relatively higher tendency to populate the periphery of the cell wall. We also calculated the amount of PFRs in each macrodomain and saw that Ori has the higher amount of PFRs while Ter has the least. This can mean that the most vital genes, which may be required for basic metabolism and growth of *E. coli* might be present in Ori. This could also be an indication of why replication would start from Ori since that would copy its most important set of genes first.

The predictive power of the integrative model is demonstrated by simulation of the chromosome contact probabilities at a resolution (500 bp) ten times higher than that measured by Hi-C experiments (5000 bp). The model was readily extendable to a partially replicating chromosome via incorporation of replication forks. In particular, the partially replicatiing state demonstrated that upon replication oriCs move towards the cell poles while dif remain predominantly at mid-cell. Finally the modular nature of the model was found to be handy in identifying the individual role of features. By a set of control simulations where the information regarding either chromosome contacts, plectonemes or both were removed from the model, we dissected the role of plectonemes and intra-chromosome contacts on its organization. We observed a greater extent of the role of chromosome contacts in its compression than plectonemes. However both plectonemes and chromosome contacts played an important role in macrodomain segregation. It should be noted that these situations presented via our control simulations would be very difficult to realize experimentally but can be very easily explored through our model.

Prior to the current initiative, there have been several attempts to model the bacterial chromosome.^11,40,51,54–56^ A previous attempt from our group had demonstrated the development of 5000bp resolution chromosome model via considering Hi-C derived data.^40,57^ A higher resolution model has also been previously proposed^51^ where the bacterial chromosome was modeled at 500bp and then back-mapped to base pair resolution. The model included plectonemes as part of the chromosome architecture which were modeled with the help of RNAP-ChIP data.^22^ But this model’s prediction of genome contact probability significantly deviates from that of Hi-C experiments. In this respect, a multi-resolution model for the bacterial chromosome that can has information about both the plectoenemes and chromosome contacts has remained wanted, which has now been reconciled in the present initiative.

However, we believe that as the model estimates 500 base pairs as 1 bead which limits one to explore phenomenon such as binding of Nucleoid Associated Proteins (NAPs) to DNA and how they influence the chromosome conformation. Thus as future directions, a more fine-grained model is on the card. Incorporation of different proteins into such a model via proper optimization of DNA - protein interactions can finally lead to a more complete model which will be more robust, accurate and provide a better understanding of the complex phenomenon of organization of the ∼4×10^6^ bp long bacterial chromosome into a ∼1*µm*^3^ cell. Finally the role of cytoplasmic particles especially ribosomes in spatial organization of the bacterial chromosome is an aspect which is currently being explored as a separate investigation.

## 5 Code Availability

We have uploaded the codes needed to generate the chromosome conformations from RNA-Seq data in Github. Please follow the link (https://github.com/JMLab-tifrh/ecoli_finer) to visit our Github repository. The modified version of the GROMACS has also been uploaded to another repository (https://github.com/JMLab-tifrh/gromacs_spherocylinder). Further data can be made available upon request.

## 6 Acknowledgment

This work was supported by computing resources obtained from shared facility of TIFR Centre for Interdisciplinary Sciences, India. We acknowledge support of the Department of Atomic Energy, Government of India, under Project Identification No. RTI 4007. JM acknowledges Core Research grants provided by the Department of Science and Technology (DST) of India (CRG/2019/001219). We would like to thank Dr. Virginia Lioy for sharing with us the cell size and the Ori/Ter ratio data. We would like to thank Dr. Mohan Joshi for insightful discussions.

## References

(1) Blattner, F. R. et al. The Complete Genome Sequence of ¡i¿Escherichia coli¡/i¿ K-12. Science 1997, 277, 1453–1462.

(2) https://bionumbers.hms.harvard.edu/bionumber.aspx?id=108589&ver=3&trm=dna,

(3) Saberi, S.; Emberly, E. Chromosome driven spatial patterning of proteins in bacteria. PLoS computational biology 2010, 6, e1000986.

(4) Verma, S. C.; Qian, Z.; Adhya, S. L. Architecture of the Escherichia coli nucleoid. PLoS genetics 2019, 15, e1008456.

(5) Holmes, V. F.; Cozzarelli, N. R. Closing the ring: links between SMC proteins and chromosome partitioning, condensation, and supercoiling. Proceedings of the National Academy of Sciences 2000, 97, 1322–1324.

(6) Dame, R. T. The role of nucleoid-associated proteins in the organization and compaction of bacterial chromatin. Molecular microbiology 2005, 56, 858–870.

(7) De Vries, R. DNA condensation in bacteria: Interplay between macromolecular crowding and nucleoid proteins. Biochimie 2010, 92, 1715–1721.

(8) Hammel, M.; Amlanjyoti, D.; Reyes, F. E.; Chen, J.-H.; Parpana, R.; Tang, H. Y.; Larabell, C. A.; Tainer, J. A.; Adhya, S. HU multimerization shift controls nucleoid compaction. Science advances 2016, 2, e1600650.

(9) Amemiya, H. M.; Schroeder, J.; Freddolino, P. L. Nucleoid-associated proteins shape chromatin structure and transcriptional regulation across the bacterial kingdom. Transcription 2021, 12, 182–218.

(10) Jun, S.; Mulder, B. Entropy-driven spatial organization of highly confined polymers: lessons for the bacterial chromosome. Proceedings of the National Academy of Sciences 2006, 103, 12388–12393.

(11) Mondal, J.; Bratton, B. P.; Li, Y.; Yethiraj, A.; Weisshaar, J. C. Entropy-based mechanism of ribosome-nucleoid segregation in E. coli cells. Biophysical journal 2011, 100, 2605–2613.

(12) Bakshi, S.; Siryaporn, A.; Goulian, M.; Weisshaar, J. C. Superresolution imaging of ribosomes and RNA polymerase in live Escherichia coli cells. Molecular microbiology 2012, 85, 21–38.

(13) Dekker, J.; Rippe, K.; Dekker, M.; Kleckner, N. Capturing chromosome conformation. science 2002, 295, 1306–1311.

(14) others,, et al. Chromosome Conformation Capture Carbon Copy (5C): a massively parallel solution for mapping interactions between genomic elements. Genome research 2006, 16, 1299–1309.

(15) others,, et al. Comprehensive mapping of long-range interactions reveals folding principles of the human genome. science 2009, 326, 289–293.

(16) Cagliero, C.; Grand, R. S.; Jones, M. B.; Jin, D. J.; OÄôSullivan, J. M. Genome conformation capture reveals that the Escherichia coli chromosome is organized by replication and transcription. Nucleic acids research 2013, 41, 6058–6071.

(17) Lioy, V. S.; Cournac, A.; Marbouty, M.; Duigou, S.; Mozziconacci, J.; Espéli, O.; Boccard, F.; Koszul, R. Multiscale structuring of the E. coli chromosome by nucleoid-associated and condensin proteins. Cell 2018, 172, 771–783.

(18) Espeli, O.; Mercier, R.; Boccard, F. DNA dynamics vary according to macrodomain topography in the E. coli chromosome. Molecular microbiology 2008, 68, 1418–1427.

(19) Wiggins, P. A.; Cheveralls, K. C.; Martin, J. S.; Lintner, R.; Kondev, J. Strong intranucleoid interactions organize the Escherichia coli chromosome into a nucleoid filament. Proceedings of the National Academy of Sciences 2010, 107, 4991–4995.

(20) Valens, M.; Penaud, S.; Rossignol, M.; Cornet, F.; Boccard, F. Macrodomain organization of the Escherichia coli chromosome. The EMBO journal 2004, 23, 4330–4341.

(21) Possoz, C.; Junier, I.; Espeli, O. wBacterial chromosome segregation. Frontiers in Bioscience-Landmark 2012, 17, 1020–1034.

(22) Grainger, D. C.; Hurd, D.; Goldberg, M. D.; Busby, S. J. Association of nucleoid proteins with coding and non-coding segments of the Escherichia coli genome. Nucleic acids research 2006, 34, 4642–4652.

(23) Scholz, S. A.; Diao, R.; Wolfe, M. B.; Fivenson, E. M.; Lin, X. N.; Freddolino, P. L. High-resolution mapping of the Escherichia coli chromosome reveals positions of high and low transcription. Cell systems 2019, 8, 212–225.

(24) Robustelli, P.; Piana, S.; Shaw, D. E. Developing a molecular dynamics force field for both folded and disordered protein states. Proceedings of the National Academy of Sciences 2018, 115, E4758–E4766.

(25) others,, et al. Free energy landscapes from SARS-CoV-2 spike glycoprotein simulations suggest that rbd opening can be modulated via interactions in an allosteric pocket. Journal of the American Chemical Society 2021, 143, 11349–11360.

(26) others,, et al. A structural model of a Ras–Raf signalosome. Nature structural & molecular biology 2021, 28, 847–857.

(27) Marrink, S. J.; Risselada, H. J.; Yefimov, S.; Tieleman, D. P.; De Vries, A. H. The MARTINI force field: coarse grained model for biomolecular simulations. The journal of physical chemistry B 2007, 111, 7812–7824.

(28) Monticelli, L.; Kandasamy, S. K.; Periole, X.; Larson, R. G.; Tieleman, D. P.; Marrink, S.-J. The MARTINI coarse-grained force field: extension to proteins. Journal of chemical theory and computation 2008, 4, 819–834.

(29) Uusitalo, J. J.; IngoÃÅlfsson, H. I.; Akhshi, P.; Tieleman, D. P.; Marrink, S. J. Martini coarse-grained force field: extension to DNA. Journal of chemical theory and computation 2015, 11, 3932–3945.

(30) others,, et al. Martini 3: a general purpose force field for coarse-grained molecular dynamics. Nature methods 2021, 18, 382–388.

(31) others,, et al. Molecular Dynamics Simulation of an Entire Cell. Frontiers in Chemistry 11, 24.

(32) Sinden, R. R.; Pettijohn, D. E. Chromosomes in living Escherichia coli cells are segregated into domains of supercoiling. Proceedings of the National Academy of Sciences 1981, 78, 224–228.

(33) Postow, L.; Crisona, N. J.; Peter, B. J.; Hardy, C. D.; Cozzarelli, N. R. Topological challenges to DNA replication: conformations at the fork. Proceedings of the National Academy of Sciences 2001, 98, 8219–8226.

(34) Bremer, H.; Dennis, P. P. Modulation of chemical composition and other parameters of the cell at different exponential growth rates. EcoSal Plus 2008, 3.

(35) Shen, B. A.; Landick, R. Transcription of bacterial chromatin. Journal of molecular biology 2019, 431, 4040–4066.

(36) Dorman, C. DNA supercoiling and transcription in bacteria: a two-way street. BMC Molecular and Cell Biology 2019, 20.

(37) Lyubchenko, Y. L.; Shlyakhtenko, L. S.; Aki, T.; Adhya, S. Atomic force microscopic demonstration of DNA looping by GalR and HU. Nucleic acids research 1997, 25, 873–876.

(38) Mäkelä, J.; Sherratt, D. J. Organization of the Escherichia coli chromosome by a Muk-BEF axial core. Molecular Cell 2020, 78, 250–260.

(39) Plunkett G. 3rd, R. K., Richardson A.J. Unpublished 2013,

(40) Wasim, A.; Gupta, A.; Mondal, J. A Hi–C data-integrated model elucidates E. coli chromosomeÄôs multiscale organization at various replication stages. Nucleic acids research 2021, 49, 3077–3091.

(41) https://bionumbers.hms.harvard.edu/bionumber.aspx?&id=101441&ver=9,

(42) Mohapatra, S.; Weisshaar, J. C. Functional mapping of the E. coli translational machinery using single-molecule tracking. Molecular microbiology 2018, 110, 262–282.

(43) Van Der Spoel, D.; Lindahl, E.; Hess, B.; Groenhof, G.; Mark, A. E.; Berendsen, H. J. GROMACS: fast, flexible, and free. Journal of computational chemistry 2005, 26, 1701–1718.

(44) Abraham, M. J.; Murtola, T.; Schulz, R.; Páll, S.; Smith, J. C.; Hess, B.; Lindahl, E. GROMACS: High performance molecular simulations through multi-level parallelism from laptops to supercomputers. SoftwareX 2015, 1, 19–25.

(45) Boles, T. C.; White, J. H.; Cozzarelli, N. R. Structure of plectonemically supercoiled DNA. Journal of molecular biology 1990, 213, 931–951.

(46) https://bionumbers.hms.harvard.edu/files/Nucleic%20Acids_Sizes_and_Molecular_Weights_2pgs.pdf,

(47) others,, et al. The three-dimensional architecture of a bacterial genome and its alteration by genetic perturbation. Molecular cell 2011, 44, 252–264.

(48) Le, T. B.; Imakaev, M. V.; Mirny, L. A.; Laub, M. T. High-resolution mapping of the spatial organization of a bacterial chromosome. Science 2013, 342, 731–734.

(49) Fisher, J. K.; Bourniquel, A.; Witz, G.; Weiner, B.; Prentiss, M.; Kleckner, N. Four-dimensional imaging of E. coli nucleoid organization and dynamics in living cells. Cell 2013, 153, 882–895.

(50) Kleckner, N.; Fisher, J. K.; Stouf, M.; White, M. A.; Bates, D.; Witz, G. The bacterial nucleoid: nature, dynamics and sister segregation. Current opinion in microbiology 2014, 22, 127–137.

(51) Hacker, W. C.; Li, S.; Elcock, A. H. Features of genomic organization in a nucleotide-resolution molecular model of the Escherichia coli chromosome. Nucleic acids research 2017, 45, 7541–7554.

(52) Hadizadeh Yazdi, N.; Guet, C. C.; Johnson, R. C.; Marko, J. F. Variation of the folding and dynamics of the e scherichia coli chromosome with growth conditions. Molecular microbiology 2012, 86, 1318–1333.

(53) Mitra, D.; Pande, S.; Chatterji, A. Polymer architecture orchestrates the segregation and spatial organization of replicating E. coli chromosomes in slow growth. Soft Matter 2022,

(54) Fritsche, M.; Li, S.; Heermann, D. W.; Wiggins, P. A. A model for Escherichia coli chromosome packaging supports transcription factor-induced DNA domain formation. Nucleic acids research 2012, 40, 972–980.

(55) Dorier, J.; Stasiak, A. Modelling of crowded polymers elucidate effects of double-strand breaks in topological domains of bacterial chromosomes. Nucleic acids research 2013, 41, 6808–6815.

(56) Chaudhuri, D.; Mulder, B. M. Bacterial Chromatin; Springer, 2018; pp 403–415.

(57) Wasim, A.; Gupta, A.; Bera, P.; Mondal, J. Interpretation of organizational role of proteins on E. coli nucleoid via Hi-C integrated model. Biophysical Journal 2022, 122, 63–81.

