## Supplementary material for "Development of a Data-driven Integrative Model of Bacterial Chromosome": SI method and figures and tables

### Methods

#### Calculation of size of beads and the simulation length scale

Let the packing fraction of DNA inside the cell be $f$. Let the length of the bacterial cell including the end caps be $L$ and the diameter be $d$. Let $l = L - d$. Therefore volume of the cell is given by

$$V_{cell} = \frac{\pi d^2}{2}(\frac{d}{3} + \frac{l}{2}) \tag{1}$$

Let the size of each bead be $\sigma$. Let there be $N$ beads where $N = \frac{4.64 \times 10^6}{res}$ where $res$ is the resolution of the model in base pairs. The volume of each bead is given by

$$V_{bead} = \frac{\pi \sigma^3}{6} \tag{2}$$

Therefore, $f \times V_{cell} = N \times V_{bead}$. Thus the size of each bead ($\sigma$) is given by

$$\sigma = d\sqrt[3]{\frac{1}{N}(1 + \frac{3l}{2d})} \tag{3}$$

Using $f = 0.1$, $res = 500$ bp, $L = 3.05$ $\mu m$, $d = 0.84$ $\mu m$, $\sigma = 31.61$ nm. However to keep simulation boxes similar to our previous studies,[1,2] we have used the simulation length scales ($\sigma_0$) as 68.1 nm. Therefore $\sigma = 0.464\ \sigma_0$, $L = 44.71\ \sigma_0$ and $d = 12.32\ \sigma_0$.

#### Force-field for the chromosome

The mass of each DNA bead has been set to $1m_0$ which is equal to 0.324 MDa[3] Therefore the complete force field for the chromosome is given by

$$V(r) = V_{adj}(r_{ij}) + V_{ex}(r_{ij}) + V_{wall}(\vec{r_i}) + V_{Hi-C}(r_{ij}) \tag{4}$$

$$V_{adj}(r_{ij}) = \frac{1}{2}k_{adj}(|\vec{r_i} - \vec{r_j}| - \sigma)^2 \tag{5}$$

where $V_{adj}(r_{ij})$ is the potential that binds adjacent beads of the polymer and $k_{adj}$ is the force constant of the springs connecting adjacent beads of the polymer. $k_{adj} = 300\ k_BT\sigma_0^{-2}$ has been used.

$$V_{ex}(r_{ij}) = \begin{cases} \frac{A}{r_{ij}^{12}} & \text{if } r_{ij} < 2^{1/6} \\ 0 & \text{otherwise} \end{cases} \tag{6}$$

where $V_{ex}(r_{ij})$ is the volume exclusion potential that prevents beads of the polymers from overlapping. We have used $A = 4.0\ k_BT\sigma_0^{12}$ for our model.

$$V_{wall}(\vec{r_i}) = \begin{cases} \frac{1}{2}k_{wall}|\vec{r_i} - \vec{R_0}|^2 & \text{if particle goes outside the confinement} \\ 0 & \text{otherwise} \end{cases} \tag{7}$$

To mimic the cell wall of the bacteria, we have used a capsule shaped confining potential $V_{wall}(\vec{r_i})$. It acts only when a particle goes outside the potential and forces the particle back towards the centre of the confinement which is given by $\vec{R_0}$. $k_{wall} = 310\ k_BT\sigma_0^{-2}$ has been used for this study.

$$V_{Hi-C}(r_{ij}) = \frac{1}{2}k_{ij}(r_{ij} - d_{ij}^{0})^2 \quad (8)$$

where $V_{Hi-C}(r_{ij})$ is the potential that maintains proper distances among the Hi-C beads obtained from experimental contact probability matrix and $k_{ij} = k_0 \exp[\frac{-(d_{ij}^{(0)} - \sigma_0)^2}{w}]$. $k_0 = 10\ k_B T \sigma_0^{-2}$ has been used for this study.

$w$ in Eq-8 needs to be optimized for proper reproducibility of the contact probability matrix from simulations. We have obtained an optimum value of $w = 1.0$. The comparison between the simulated and experimental contact probability matrices have been shown in Figure 1.

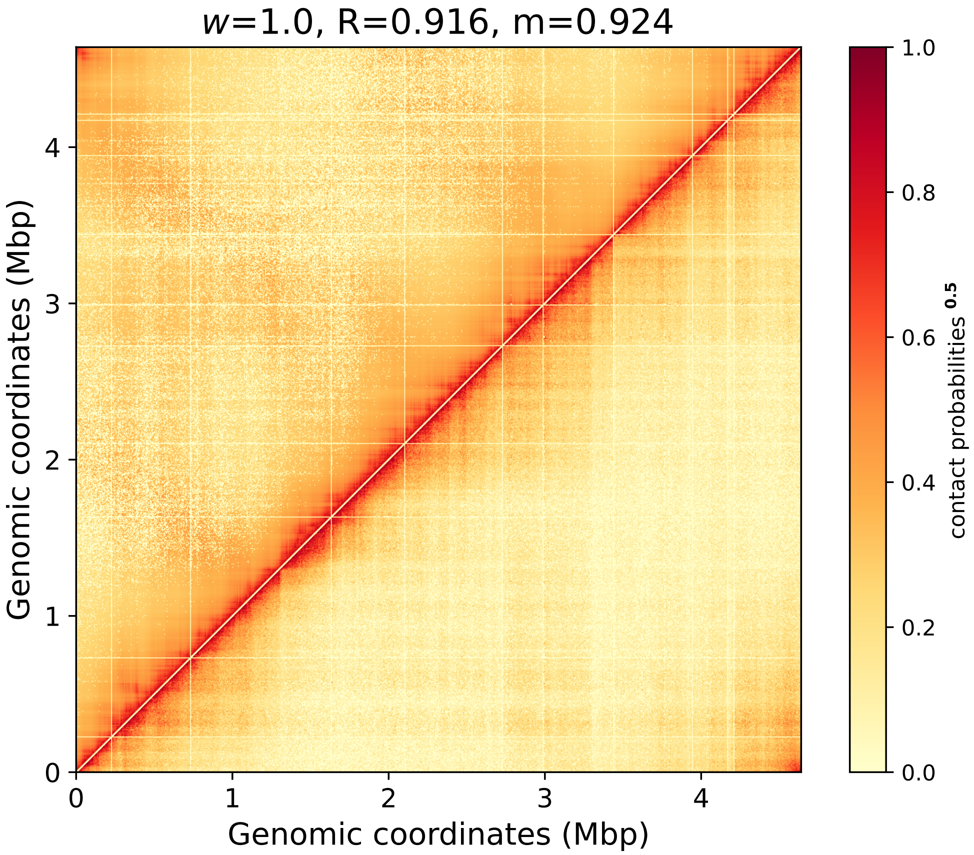


Figure 1: Comparison of experimental and simulated contact probability matrix for $w = 1.0$. Upper triangle represents probabilities obtained via simulations while the lower triangle denotes the experimental matrix. The values have been enhanced (by using $\sqrt{\text{contact probabilities}}$) for a better visual comparison.

#### Modeling ribosomes and polysomes

Ribosomes comprise of 30S and 50S sub-units in *E. coli.* The 30S and 50S together can form a 70S ribosome complex, which polymerizes to form polysomes. Polysomes are the function form of ribosomes that help in translation of mRNA into proteins. Each 30S, 50S and 70S monomeric subunit have been modeled as spherical particles with masses and sizes (in terms of $\sigma_0$) as provided in Table-S1. The polysomes have been modeled as a 13-mer of 70S subunits.[4] We have incorporated 21000 particles belonging to 30S, 50S and 70S. Out of them 1292 are 30S, 1292 are 50S and 18416 particles (=2102 polysome chains) are 70S.[5]

#### Force-field for the ribosomes and polysomes

The ribosomes and polysomes have interactions with themselves, with the DNA and with the wall ($V_{wall}(\vec{r_i})$). The $V_{wall}$ confines ribosomes and polysomes in the cell wall and has the same form as Eq 7.

The ribosomes and polysomes with themselves have purely repulsive interactions which has the same form as Eq 6. The value of $A$ between ribosomes have been provided in Table-S2.

The monomeric beads of the polysomes are tightly bound together via a harmonic potential which has the same form as Eq 5 and given by

$$V_{adj}^{polysome}(r_{ij}) = \frac{1}{2}k_{adj}^{polysome}(|\vec{r_i} - \vec{r_j}| - 0.307\sigma_0)^2 \tag{9}$$

where $k_{adj}^{polysome} = 17000\ k_B T \sigma_0^{-2}$.

#### Non-bonded interactions among particles

DNA-DNA, ribosome-ribosome, polysome-polysome and ribosome-polysome interactions are purely repulsive and have the same form as of Eq 6. Values of $A$ for respective pairs have been provided in Table-S2.

However, complete Lennard-Jones potential has been considered for the ribosomal or polysomal interaction with DNA and is given by

$$V_{DNA-rib/poly}(r_{ij}) = \frac{c_{12}}{r_{ij}^{12}} - \frac{c_6}{r_{ij}^{6}} \tag{10}$$

where $c_6$ and $c_{12}$ have been tuned to match the simulated and experimental linear density profiles of DNA and ribosomes.

Details of $c_6$ and $c_{12}$ have been provided in Table-S2.

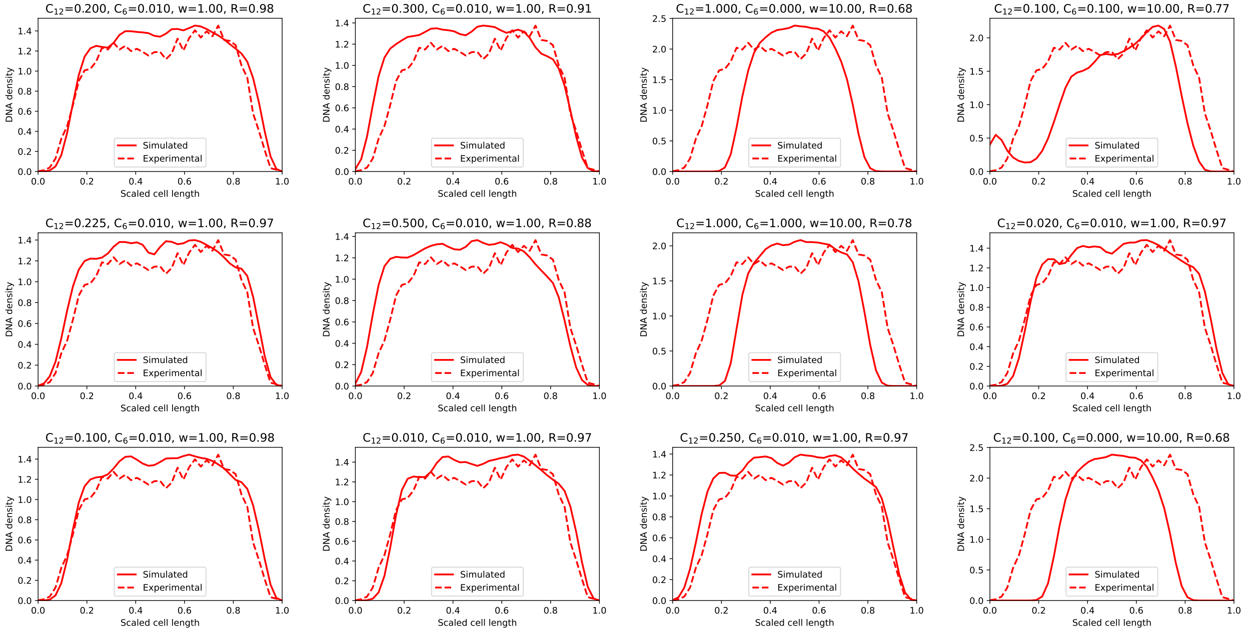


Figure 2: The DNA densities for different interaction parameter sets.

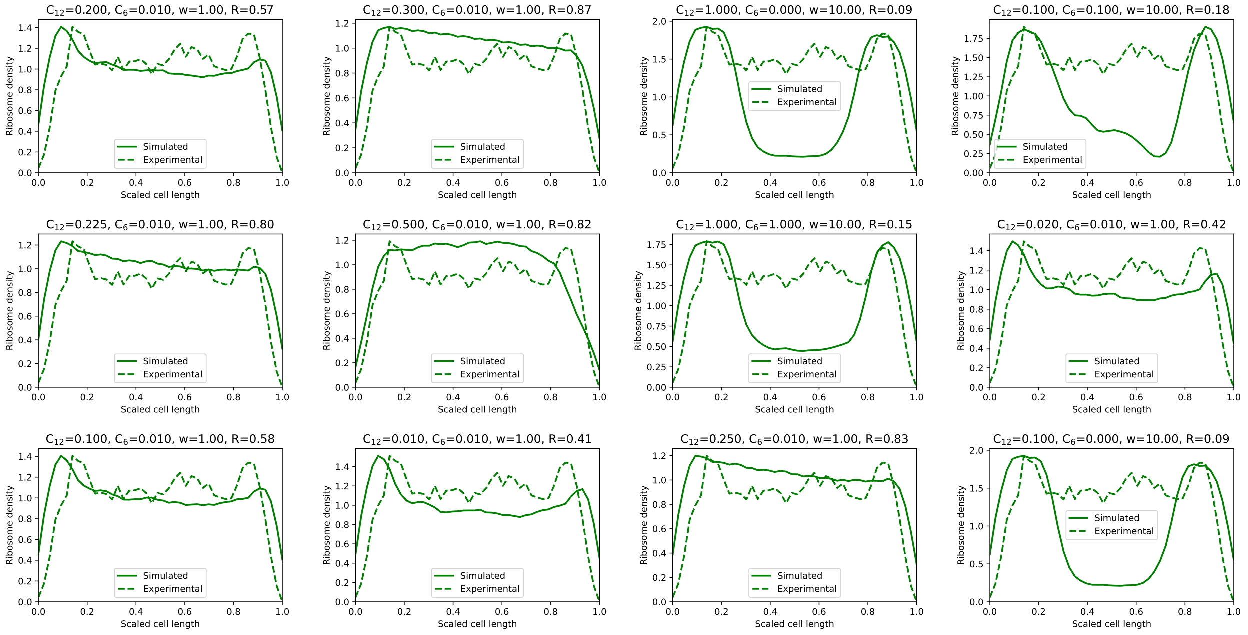


Figure 3: The DNA densities for different interaction parameter sets.

#### Interaction of probes with the particles in the system

Let the size of the probe be $\sigma_{probe}$, size of a 30S ribosome be $\sigma_{30S}$, size of a 50S ribosome be $\sigma_{50S}$, size of a 70S ribosome be $\sigma_{70S}$ and size of a bead of the polymer representing the chromosome be $\sigma_{DNA}$. Except for $\sigma_{probe}$, all other sizes have been provided in Table-S1. The interaction of a probe with itself and any other particle is purely repulsive and is given by $V_{probe}(r) = \frac{A_{probe}}{r^{12}}$ where $A_{probe} = 4 * \sigma'^{12}$ and $\sigma'$ is the effective size of the particles given by $\sigma' = \sqrt{\sigma_{probe}\sigma_{30S/50S/70S/DNA}}$. We have used probe sizes of 25 nm, 40 nm, 45 nm 47.5 nm, 50 nm, 52.5 nm, 55 nm and 60 nm.

Therefore if we wanted to calculate the interaction potential between a probe and a 30S ribosome particle, then $\sigma' = \sqrt{\sigma_{probe} * \sigma_{30S}}$ and $V_{probe/30S}(r) = \frac{4\sigma'^{12}}{r^{12}}$. If the probe size is 50 nm, $\sigma_{probe} = \frac{50}{68.14} = 0.733\sigma_0$ and $\sigma' = 0.387\sigma_0$. The corresponding
$A_{probe} = 4 * (0.387\sigma_0)^{12} = 4.6 \times 10^{-5}\sigma_0^{12}$.

### Supplementary Tables

Table S1: Parameters for ribsosomes

| particle type | mass ($m_0 = 0.324$ MDa) | size ($\sigma_0 = 68.14$ nm) |
|---|---|---|
| DNA | 1 | 0.464 |
| 30S ribsosome | 2.62 | 0.205 |
| 50S ribsosome | 4.19 | 0.249 |
| 70S ribsosome | 7.12 | 0.307 |

Table S2: All non-bonded interaction parameters.

| particle-1 | particle-2 | $c_6$ ($k_BT\sigma_0^6$) | $A/c_{12}$ ($k_BT\sigma_0^{12}$)** |
|---|---|---|---|
| DNA | DNA | —— | 1.0 |
| 30S ribosomes | 30S ribosomes | —— | $2.236\times10^{-8}$ |
| 50S ribosomes | 50S ribosomes | —— | $2.298\times10^{-7}$ |
| 70S ribosomes | 70S ribosomes | —— | $2.901\times10^{-6}$ |
| 30S ribosomes | 50S ribosomes | —— | $7.167\times10^{-8}$ |
| 50S ribosomes | 70S ribosomes | —— | $8.164\times10^{-7}$ |
| 30S ribosomes | 70S ribosomes | —— | $2.547\times10^{-7}$ |
| DNA | 30S ribosomes | $3.459\times10^{-4}$ | $2.990\times10^{-8}$ |
| DNA | 50S ribosomes | $3.459\times10^{-4}$ | $9.587\times10^{-8}$ |
| DNA | 70S ribosomes | $3.459\times10^{-4}$ | $3.406\times10^{-8}$ |

**For same particle and all ribosome-ribosome interactions, there no $c_6$ and only $A$. For DNA-ribosome interactions only should consider $c_6$ and $c_{12}$.

#### Supplementary Figures

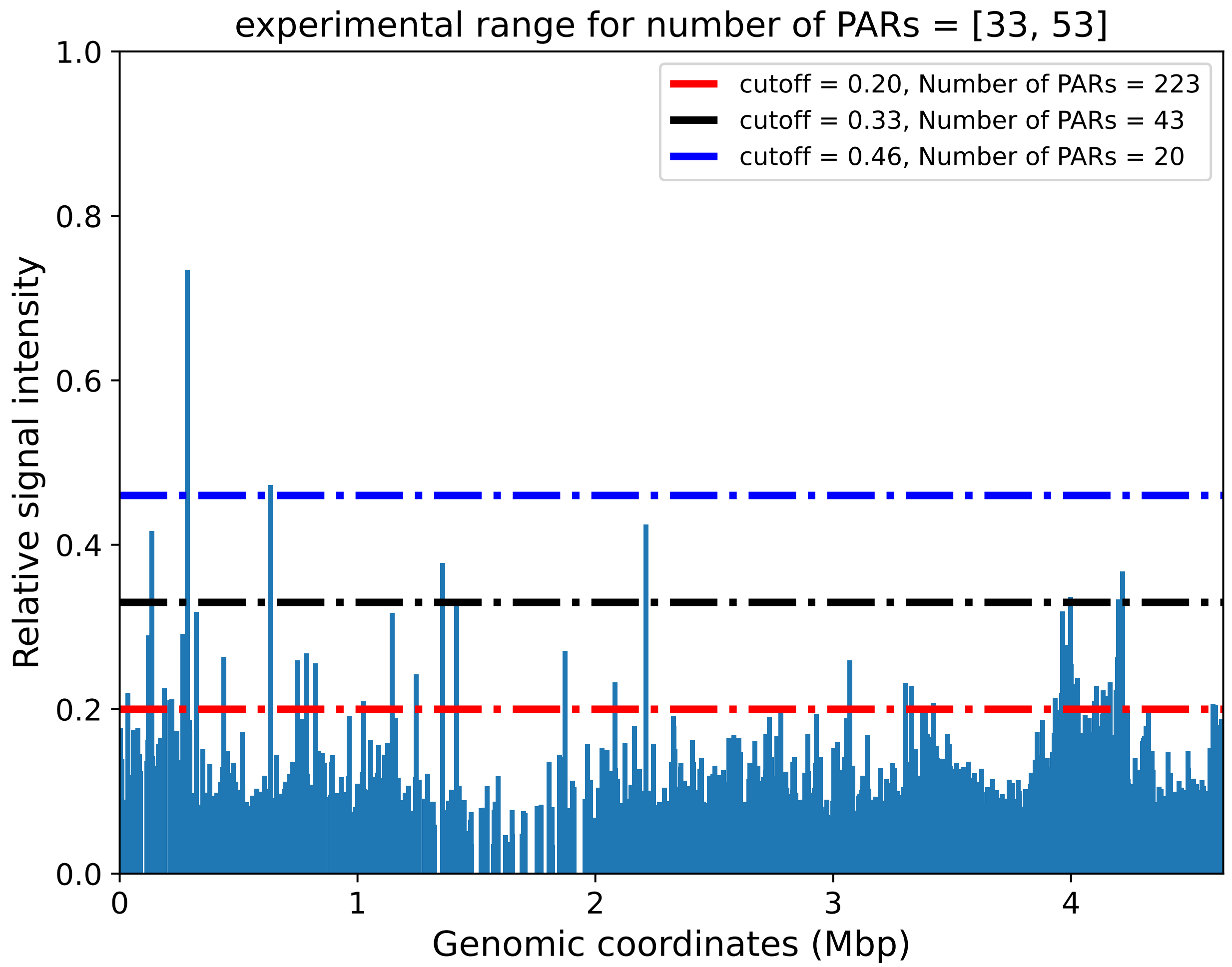


Figure S1: The figure shows that the cutoff obtained is optimum for the number of PARs present in the system. The optimum cutoff = 0.33.

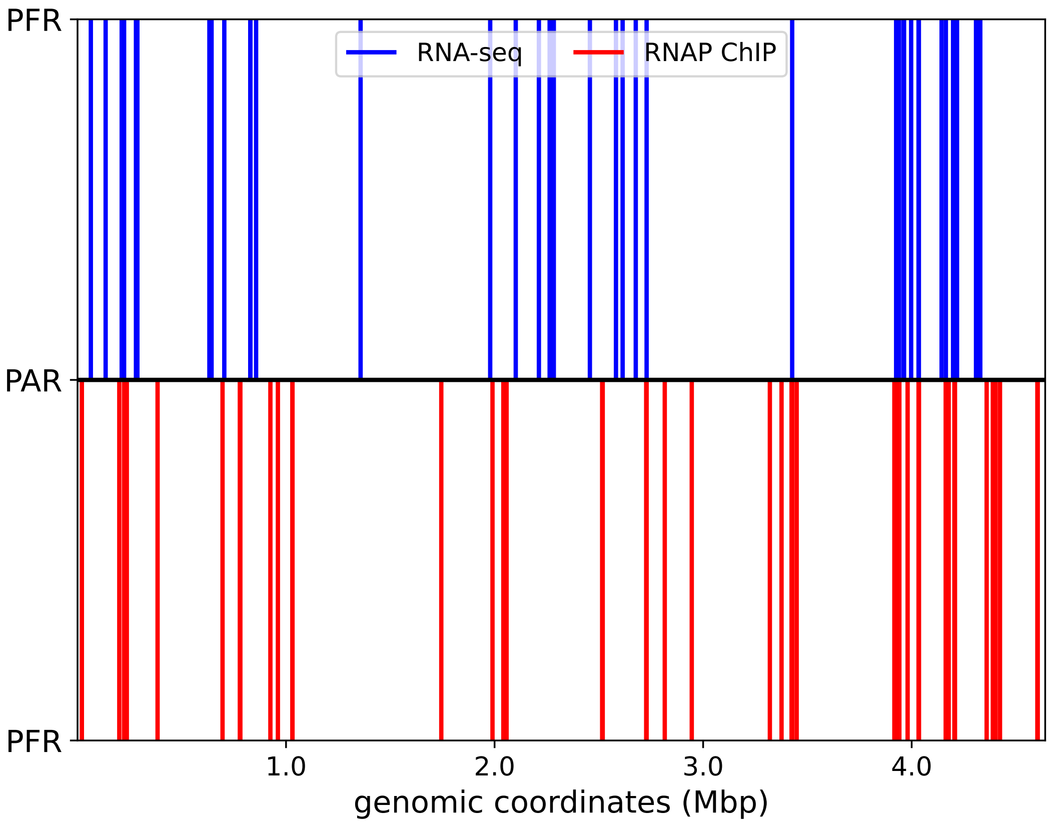


Figure S2: Here the blue and the red vertical lines represent the plectoneme free regions (PFRs) as obtained from RNA-Seq and RNAP-ChIP data, respectively. From the figure we can observe a very high degree of similarity in the PFRs predicted using our protocol for both datasets.

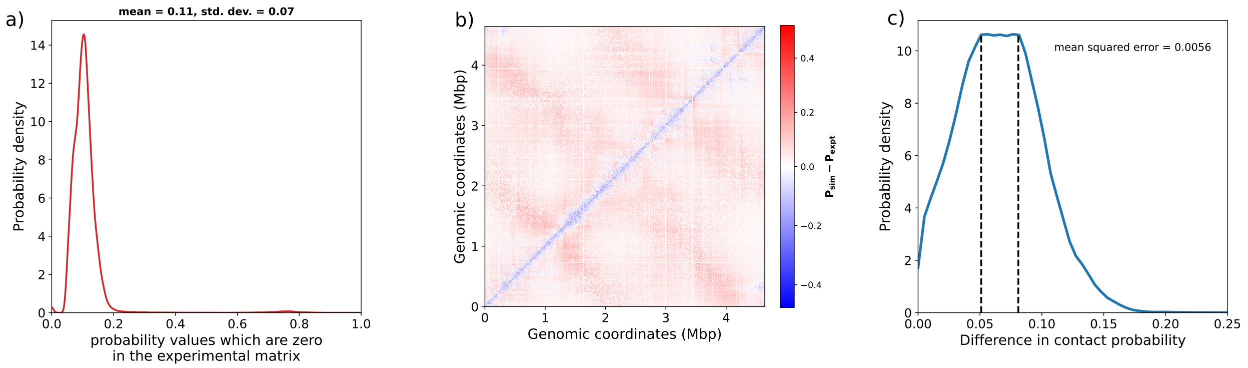


Figure S3: **a)** Distribution of the probabilities which were filtered out.. **b)** Difference matrix between simulated and experimental contact probabilities. **c)** Distribution of absolute differences. The dashed lines represent the plateau region of the maxima.

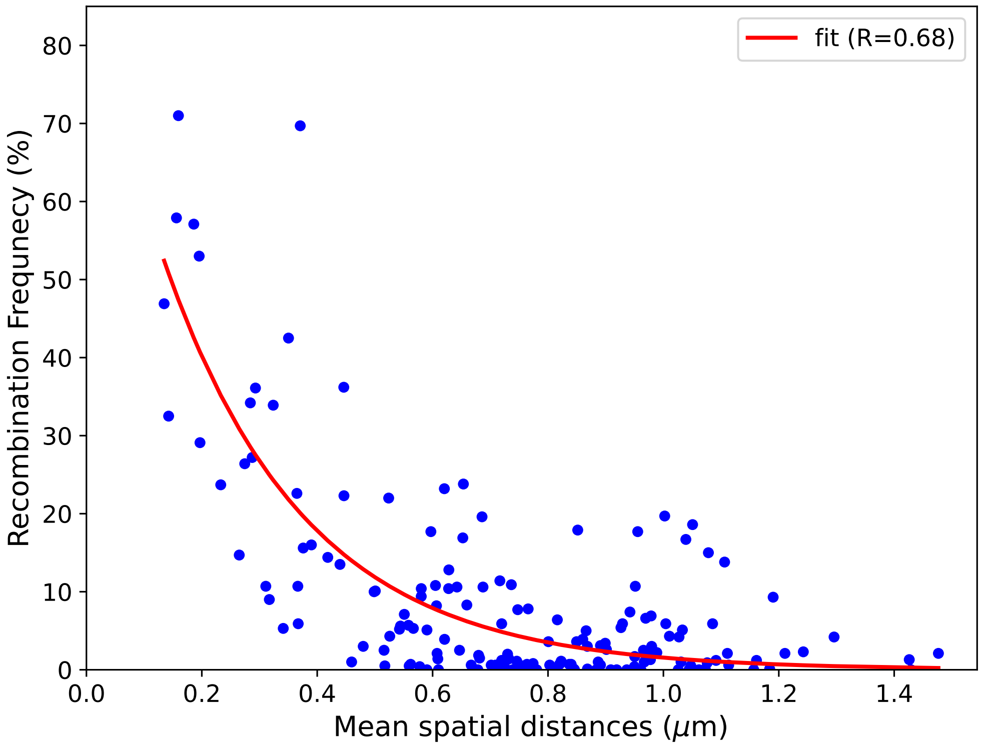


Figure S4: The mean spatial distance along pairs of probes were obtained from the simulations and plotted along the x-axis. The recombination frequencies of the respective probe pairs were plotted along the y-axis. The obtained points were fitted with an exponentially decaying function.

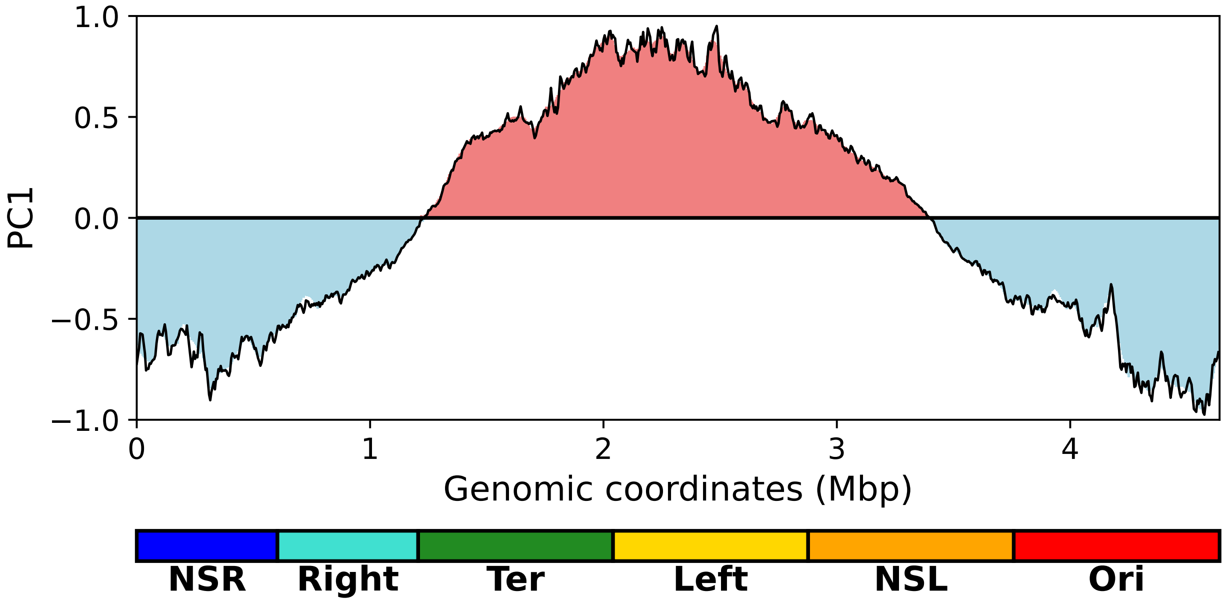


Figure S5: PC1 vs genomic coordinates for the experimental contact probability matrix at 5 Kbp. Red regions indicate positive values while blue regions indicate negative values. Macrodomains have been marked below the x-axis.

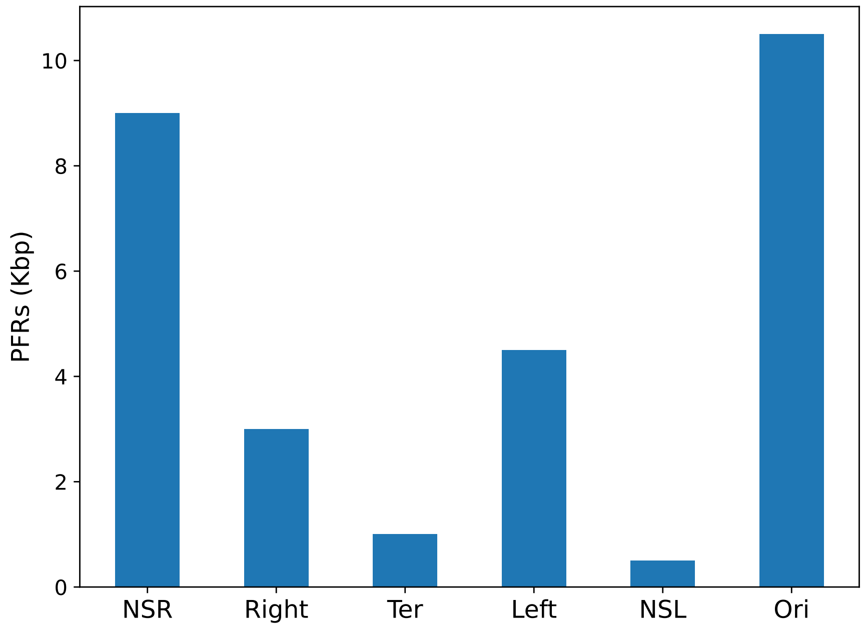


Figure S6: PFR content of each macrodomain.

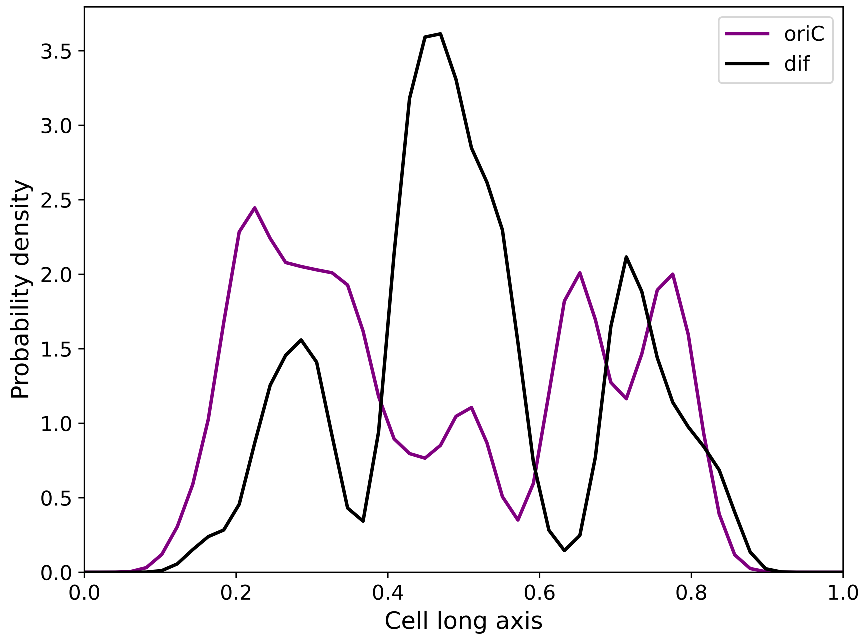


Figure S7: oriC and dif position distribution

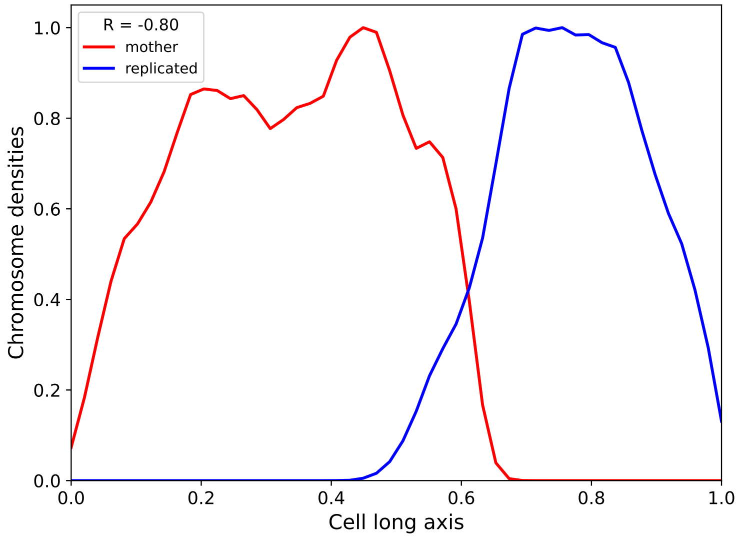


Figure S8: Linear densities along the long axis of the cell for the mother chromosome (red) and replicated chromosome (blue). The pearson correlation coefficient (R) is -0.8 which suggests proper segregation of the mother and replicated chromosomes.

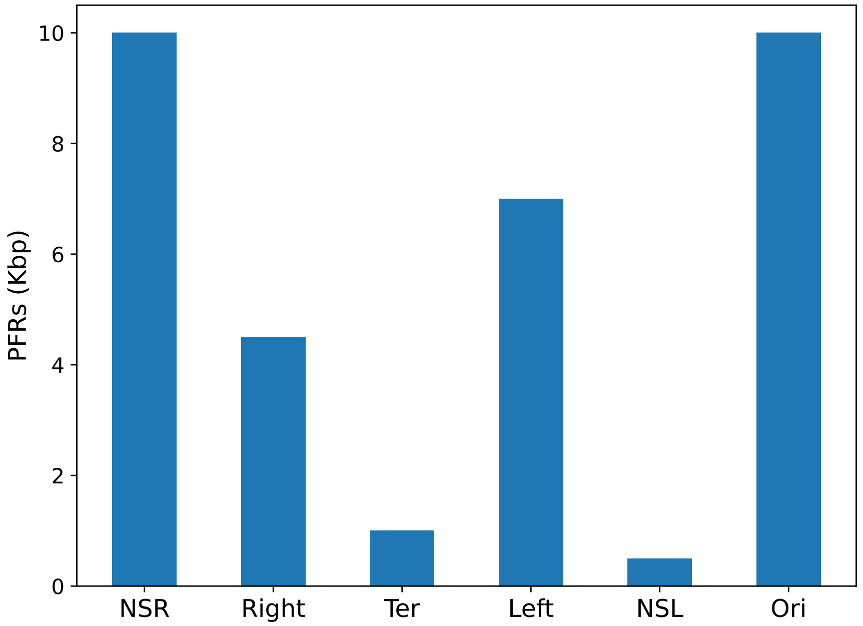


Figure S9: PFR content of each macrodomain for G=1.6.

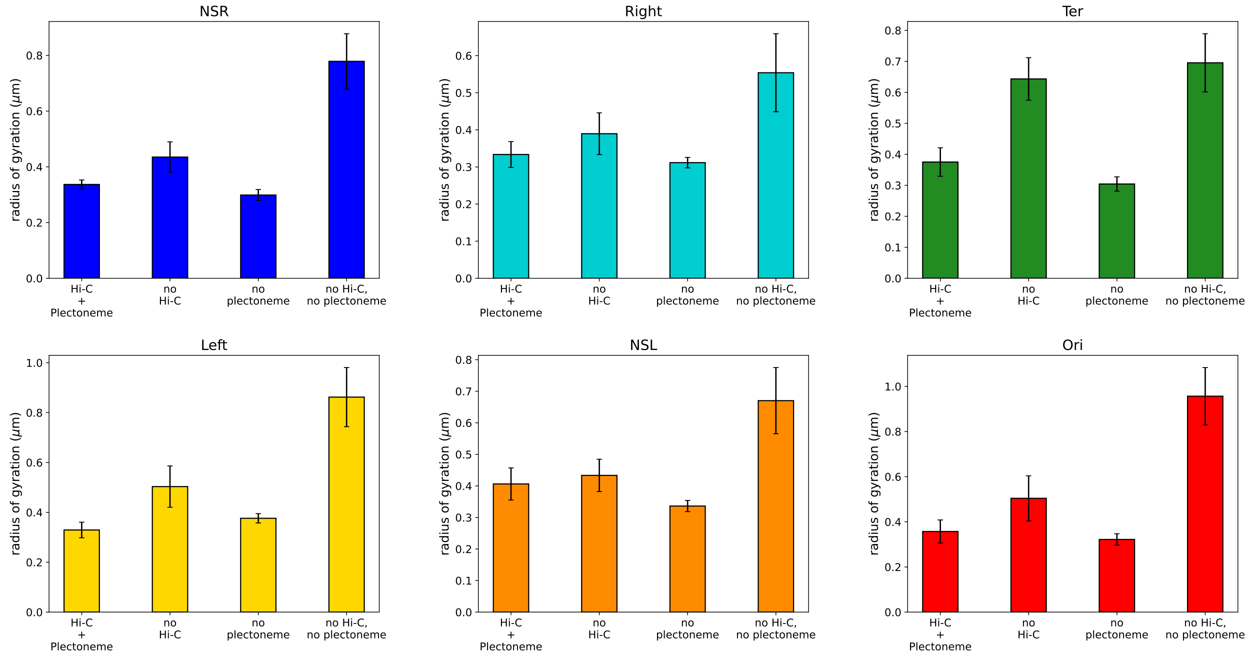


Figure S10: Radius of gyration (a measure of size) for each macrodomain for each control scenario.

### References


(1) Abdul Wasim, Ankit Gupta, and Jagannath Mondal. A hi–c data-integrated model elucidates e. coli chromosomes multiscale organization at various replication stages. *Nucleic acids research*, 49(6):3077–3091, 2021. 1

(2) Palash Bera, Abdul Wasim, and Jagannath Mondal. Hi-c embedded polymer model of escherichia coli reveals the origin of heterogeneous subdiffusion in chromosomal loci. *Physical Review E*, 105(6):064402, 2022. 1

(3) `https://bionumbers.hms.harvard.edu/files/Nucleic%20Acids_Sizes_and_Molecular_Weights_2pgs.pdf`. 2

(4) Jagannath Mondal, Benjamin P Bratton, Yijie Li, Arun Yethiraj, and James C Weisshaar. Entropy-based mechanism of ribosome-nucleoid segregation in e. coli cells. *Biophysical journal*, 100(11):2605–2613, 2011. 4

(5) Sonisilpa Mohapatra and James C Weisshaar. Functional mapping of the e. coli transla-

tional machinery using single-molecule tracking. *Molecular microbiology*, 110(2):262–282, 2018. 4
